# Mitochondria-targeted hydrogen sulfide donor reduces fatty liver and obesity in mice fed a high fat diet by inhibiting *de novo* lipogenesis and inflammation *via* mTOR/SREBP-1 and NF-κB signaling pathways

**DOI:** 10.1101/2024.04.17.589169

**Authors:** Aneta Stachowicz, Klaudia Czepiel, Anna Wiśniewska, Kamila Stachyra, Magdalena Ulatowska-Białas, Beata Kuśnierz-Cabala, Marcin Surmiak, Grzegorz Majka, Katarzyna Kuś, Mark E. Wood, Roberta Torregrossa, Matthew Whiteman, Rafał Olszanecki

## Abstract

**Background:** Metabolic diseases that include obesity and metabolic-associated fatty liver disease (MAFLD) are a rapidly growing worldwide public health problem with unmet clinical need. The pathogenesis of MAFLD is very complex including abnormally increased lipogenesis, chronic inflammation, mitochondrial dysfunction, and oxidative stress. A growing body of evidence suggests that hydrogen sulfide (H_2_S) is an important player in the liver, impacting lipid metabolism and mitochondrial function. However, direct delivery of H_2_S to mitochondria has not been investigated as a therapeutic strategy in obesity-related metabolic disorders. Therefore, the aim of our study was to comprehensively evaluate the influence of prolonged treatment with a mitochondria sulfide delivery molecule (AP39) on the development of fatty liver and obesity in a high fat diet (HFD) fed mice.

**Results:** Our results demonstrated that AP39 reduced fatty liver in HFD-fed mice, which was corresponded with decreased triglyceride content in the liver and plasma as well as increased GSH/GSSG ratio in the plasma. Furthermore, treatment with AP39 downregulated pathways related to biosynthesis of unsaturated fatty acids, lipoprotein assembly and PPAR signaling in the liver of HFD-fed mice. It also led to a decrease in *de novo* lipogenesis in the liver by downregulating mTOR/SREBP-1/SCD1 signaling pathway. Moreover, AP39 administration alleviated obesity in HFD-fed mice, which was reflected by reduced weight of mice and adipose tissue, decreased leptin levels in the plasma and upregulated expression of ATGL, a lipolysis enzyme in epididymal white adipose tissue (eWAT). Finally, AP39 reduced inflammation in the liver and eWAT measured as the expression of pro inflammatory markers (*Il1b, Il6, Tnf*, *Mcp1*), which was due to the downregulation of mTOR/NF-κB signaling pathway.

**Conclusions:** Taken together, mitochondria-targeted sulfide delivery molecules could potentially provide a novel therapeutic approach to the treatment/prevention of obesity-related metabolic disorders.

## Introduction

Metabolic diseases that include obesity, nonalcoholic fatty liver disease (NAFLD), diabetes, hyperlipidemia etc., are currently a worldwide public health problem in the Western countries^1^. Obesity is an abnormal accumulation of white adipose tissue (WAT) associated with chronic inflammation and insulin resistance ^2^ and it is the most common risk factor for the development of NAFLD ^3^. NAFLD, recently re-termed as metabolic-associated fatty liver disease (MAFLD), is a complex common liver disorder in the world manifested by triglyceride accumulation in the cytoplasm of hepatocytes ^4^. It encompasses a wide spectrum of pathological conditions ranging from a simple hepatic steatosis, steatosis with inflammatory response - nonalcoholic steatohepatitis (NASH), cirrhosis and fibrosis, and finally hepatocarcinoma ^5^. The pathogenesis of MAFLD is very complex, including lipotoxicity, altered lipogenesis, lipolysis and fatty acid beta-oxidation, chronic inflammation, mitochondrial dysfunction, and oxidative stress ^6^. Particularly, *de novo* lipogenesis, the synthesis of new fatty acids from acetyl-CoA subunits, was shown to be abnormally increased in MAFLD and involved in its aetiology ^7^. Important transcription factor regulating the expression of lipogenic genes such as fatty acid synthase (FASN) and stearoyl-CoA desaturase 1 (SCD1), is sterol regulatory element binding protein 1 (SREBP1). Mounting evidence indicates that SREBP1 transcription factor is induced by the mechanistic (mammalian) target of rapamycin (mTOR) pathway, which activates hepatic *de novo* lipogenesis ^8^. However, numerous aspects of pathogenesis of MAFLD remain unclear and there are no effective treatment strategies available.

A growing body of evidence suggests that hydrogen sulfide (H_2_S) is an important player in the physiological and pathological processes in the liver, including lipid metabolism and fibrosis ^9^. Dysregulation of H_2_S synthesis and transmission was observed in MAFLD, diabetes, liver fibrosis and cancer, among others ^10,11^. H_2_S is a small gaseous signaling molecule that has anti-inflammatory, anti-oxidant and anti-fibrotic properties ^12,13^. It is endogenously produced by two cytosolic enzymes, cystathionine-β-synthase (CBS) and cystathionine-γ-lyase (CSE) as well as 3-mercaptopyruvate sulfurtransferase (3-MST), both a mitochondrial and cytosolic enzyme ^14^. Of note, higher concentrations of H_2_S are toxic by inhibiting complex IV of electron transport chain (ETC), ^15^ whereas low concentrations of H_2_S support cellular bioenergetics ^16^. Intramitochondrial H_2_S derived from 3-MST enzyme was shown to serve as an electron donor to complex II of ETC, thus, complementing the bioenergetic role of Krebs cycle as an alternative source of electrons for oxidative phosphorylation and ATP production ^16^. Importantly, oxidative stress may suppress this effect, which could have implications in the pathogenesis of various diseases, such as MAFLD and obesity ^17^. However, direct delivery of H_2_S to mitochondria has previously not been considered as a therapeutic strategy in the treatment of obesity and MAFLD.

Interestingly, a new slow-releasing, mitochondria-targeted, H_2_S donor AP39 has been recently synthesized. It consists of mitochondria targeting motif – triphenylphosphonium (TPP^+^) linked to H_2_S donating moiety. AP39 at lower concentrations was shown to stimulate cellular bioenergetics and reduce oxidative stress ^18,19^. Moreover, AP39 was demonstrated to protect myocardium against reperfusion injury ^20^. Therefore, the aim of our current study was to comprehensively investigate the effects of prolonged treatment with direct mitochondria-targeted H_2_S donor AP39 on the development of fatty liver and obesity in C57BL/6J mice on a high-fat diet (HFD).

## Materials and Methods

### Animal studies

Forty-eight male C57BL/6J (JAX^TM^) mice were obtained from the Jackson Laboratory (Bar Harbor, CA, USA). The animals were maintained on 12 h dark/12 h light cycles at room temperature (22.5 ± 0.5 °C) and 45-55% humidity with access to water ad libitum and diet. At the age of 8 weeks the mice were fed with a normal chow diet or an HFD (Sniff (Soest, Germany): E15744-344 containing 45% fat) for 12 weeks. The animals were divided into four groups: C57BL/6J mice treated with vehicle (10% DMSO) (control) (*n* = 12), C57BL/6J mice on an HFD treated with vehicle (10% DMSO) (HFD) (*n* = 12), control mice treated with AP39 (control + AP39) (*n* = 12) and C57BL/6J mice on an HFD treated with AP39 (HFD + AP39) (*n* = 12). AP39 was synthesized in-house as previously described by us ^21^. It was administered subcutaneously to the mice at a dose of 100 nM per kg of body weight per day three days a week for 12 weeks. At the age of 5 months the mice were euthanized 5 min after injection of Fraxiparine (Nadroparin) i.p (1000 UI; Sanofi-Synthelabo, Paris, France) in chamber filled with carbon dioxide at a rate of 20-30% CO_2_ chamber volume per minute, in accordance with AVMA Panel 2007 recommendations and institutional IACUC guidelines. The selected tissues (liver, epididymal white adipose tissue (eVAT), subcutaneous inguinal white adipose tissue (iWAT) and brown adipose tissue (BAT)) were dissected, and the blood was collected. All animal procedures were conformed with the guidelines from Directive 2010/63/EU of the European Parliament on the protection of animals used for scientific purposes and were approved by the Jagiellonian University Ethical Committee on Animal Experiments (No. 517/2021).

### Glucose tolerance test (GTT)

Glucose tolerance test (GTT) was performed 48 h before euthanization. Briefly, mice were fasted for 5 h with free access to water, followed by an oral administration of glucose (2 g/kg). The blood samples were collected from the caudal vein for the measurements of fasting blood glucose and fasting insulin levels. The blood glucose concentration was estimated using Roche ACCU-CHEK glucometer (Aviva, London, UK) at 0, 15, 30, 60, 120 min. The area under the curve (AUC) was calculated as the area between the GTT line and baseline using GraphPad Prism 9.3.1 software. The blood for insulin measurement was collected in a tube with 0.5 M EDTA and then centrifugated at 13000 × g for 1 min. Serum insulin concentration was measured by the Insulin Rodent (Mouse/Rat) Chemiluminescence ELISA (ALPCO, Salem, NH, USA), according to the manufacturer’s guidelines. The insulin resistance score (HOMA-IR) was calculated as following: fasting blood glucose (mg/dL) multiplied by fasting serum insulin (mU/L) divided by 405.

### Biochemical Measurements

The blood was centrifuged at 1000 × g at 4°C for 10 min and the plasma was collected and stored at - 80°C. The concentration of H_2_S in the liver was measured by methylene blue assay using commercially available H_2_S Colorimetric Assay Kit from Elabscience (Houston, TX, USA). The levels of total cholesterol, triglycerides (TG), low-density lipoproteins (LDL), high-density lipoproteins (HDL), aspartate aminotransferase (AST) and alanine aminotransferase (ALT) were measured using an enzymatic method on a Cobas 8000 analyzer (Roche Diagnostics, Indianapolis, IN, USA). The content of TG in the liver was assayed using the Triglyceride Colorimetric Assay Kit (Cayman Chemical, Ann Arbor, MI, USA), according to the manufacturer’s guidelines. Plasma concentrations of fatty acid binding protein 4 (FABP4) and leptin were determined using the xMAP technology Luminex assays (R&D Systems, Minneapolis, MN, USA) and the Luminex MAGPIX System (Luminex Corp., Austin, TX, USA). The ratio of reduced glutathione (GSH) to oxidized glutathione (GSSG) was measured by commercially available Glutathione Colorimetric Detection Kit (Invitrogen, Waltham, MA, USA).

### Histology

The samples of the liver, eWAT, iWAT and BAT tissues were fixed using formalin and embedded in paraffin. The paraffin sections (2 µm thickness) were stained with hematoxylin-eosin (H&E) method. Samples were assessed microscopically for the presence of microvesicular and macrovesicular steatosis. The mean percentage of steatotic hepatocytes was specified in each case. Moreover, the maintenance of lobular structure of the liver as well as the presence of both inflammatory infiltrates and necrotic changes were evaluated. For eWAT and iWAT diameters of cells were measured as follows: for each case, measurements were taken in four randomly selected areas of the slide, each time five fat cells lying next to each other were evaluated. Measurements were made under a 40x objective - at this magnification, 40 corresponds to a diameter of 0.1 mm. Furthermore, for each case, the number of clusters of inflammatory cells in the adipose tissue (number of inflammatory foci) was assessed, giving their total number in an area of 10 mm^2^. For BAT tissue, in each slide, the number of lipocyte nuclei in five randomly selected fields at magnification 40x (an area of 0.0625 mm^2^) were counted and the average number of cell nuclei was calculated for each case.

### Quantitative reverse transcription polymerase chain reaction (RT‒qPCR)

Briefly, total RNA was isolated from the liver and eWAT using the ReliaPrep^TM^ RNA Tissue Miniprep System (Promega, Madison, WI, USA) and the RNeasy Fibrous Tissue Mini Kit (Qiagen, Hilden, Germany), respectively, according to the manufacturer’s instructions. The RNA concentration of each sample was measured at a wavelength of 260 nm (A260) in a Synergy H1 microplate reader (BioTek Instruments, Inc., Winooski, VT). The purity of the extracted total RNA was determined by the A260/A280 ratio. cDNA was synthesized by the reverse transcription of 1000 ng of total RNA from each sample, using a High-Capacity Reverse Transcription Kit (Applied Biosystems, Foster City, CA). The cDNA was diluted ten times prior to PCR amplification. The primers used in our experiments included primers purchased from Bio-Rad (*Srebf1*, *Fasn*, *Cd36, Cpt1a, Pparg, Ppara*), RealTimePrimers (*Il1b, Il6, Tnf, Mcp1, Rpl13a*) and Merck Millipore (*Stx5a*) as well as following primers *Lep*: 5’ GTGGCTTTGGTCCTATCTGTC 3’ (forward), 5’ CGTGTGTGAAATGTCATTGATCC 3’ (reverse); *Atgl*: 5’ GATTGCGAAGGTTGAACTGGAT 3’ (forward), 5’ CTCAGGCGAGAGTGACATCT 3’ (reverse); *Lipe*: 5’ CGGCGGCTGTCTAATGTCT 3’ (forward), 5’ CGTTGGCTGGTGTCTCTGT 3’ (reverse); *Dgat1*: 5’ GGAATATCCCCGTGCACAA 3’ (forward), 5’ CATTTGCTGCTGCCATGTC 3’ (reverse); *Dgat2*: 5’ CCGCAAAGGCTTTGTGAA 3’ (forward), 5’ GGAATAAGTGGGAACCAGATCAG 3’ (reverse). 2x SsoAdvanced™ Universal SYBR® Green Supermix (Bio-Rad, Hercules, CA, USA) was used to carry out real-time PCR reaction. Analysis of relative gene expression was performed by the CFX96 Touch Real-Time PCR Detection System (Bio-Rad, Hercules, CA, USA) with *Stx5a* and *Rpl13a* as internal reference genes for liver and eVAT, respectively. Data were analyzed using the 2–ΔΔCt method in an Excel spreadsheet.

### Liquid chromatography-tandem MS (LC‒MS/MS) analysis of mouse liver

Mouse liver was homogenized using a Tissue Lyser LT (Qiagen, Hilden, Germany) and lysed in a buffer containing 0.1 M Tris-HCl, pH 7.6, 2% sodium dodecyl sulfate, and 50 mM dithiothreitol (Sigma Aldrich, St. Louis, MO, USA) at 96 °C for 10 min. A total protein concentration in lysates and the peptide contents in the digests were assayed using a tryptophan fluorescence based WF-assay ^22^. Seventy micrograms of protein were digested overnight using the filter-aided sample preparation (FASP) method ^23^ with Trypsin/Lys-C mix (Promega, Madison, WI, USA) (enzyme-to-protein ratio 1:35) as the digestion enzymes. Next, the samples were purified with C18 Ultra-Micro Spin Columns (Harvard Apparatus, Holliston, MA, USA). All samples were dissolved in 0.1% formic acid at a concentration of 0.5 µg/µl and spiked with the iRT peptides (Biognosys, Schlieren, Switzerland).

One microgram of peptide was injected into a nanoEase^TM^ M/Z Peptide BEH C18 75 µm i.d. × 25 cm column (Waters, Milford, MA, USA) via a nanoEase^TM^ M/Z Symmetry C18 180 µm i.d. × 2 cm trap column (Waters, Milford, MA, USA) and separated using a 1% to 40% B phase linear gradient (A phase - 0.1% FA in water; B phase - 80% ACN and 0.1% FA) with a flow rate of 250 nL/min on an UltiMate 3000 HPLC system (Thermo Scientific, Waltham, MA, USA) coupled to a Orbitrap Exploris™ 480 Mass Spectrometer (Thermo Scientific, Waltham, MA, USA). The nanoelectrospray ion source parameters were as follows: ion spray voltage: 2.2 kV, ion transfer tube 275 °C. For data-independent (DIA) acquisition, spectra were collected for 145 min in full scan mode (400–1250 Da), followed by 55 DIA scans using a variable precursor isolation window approach and AGC set to custom 1000%.

The DIA MS data were analyzed in Spectronaut (Biognosys, Schlieren, Switzerland) ^24^ software using directDIA^TM^ approach. MS data were filtered by 1% FDR at the peptide and protein levels, while quantitation was performed at the MS2 level and global imputation with a missingness rate set to 0.5 was used. Statistical analysis of differential protein abundance was performed at both the MS1 and MS2 levels ^25^ using unpaired t tests with multiple testing correction after Storey ^26^.

### Constructing protein‒protein interaction networks

Functional grouping and pathway analysis were performed using PINE (Protein Interaction Network Extractor) software ^27^ with the STRING and GeneMANIA databases using a score confidence of 0.4 and a ClueGO p value cutoff < 0.05. The mass spectrometry data have been deposited to the ProteomeXchange Consortium via the PRIDE partner repository ^28^ with the dataset identifier PXD051150.

### Western blot analysis

Mouse liver and eVAT were lysed in 2% SDS, 50 mM DTT in 0.1 M Tris-HCl (pH 7.6) as described in proteomics section. Tissue lysates containing equal amounts of total protein were mixed with 4x Laemmli Sample Buffer (Bio-Rad, Hercules, CA, USA) and incubated at 96 °C for 5 min. Protein samples (10 µg of protein per well) were separated on SDS-polyacrylamide gels (Criterion^TM^ TGX^TM^ Precast Gels, 4-15%; Bio-Rad, Hercules, CA, USA) using a Laemmli buffer system and then semidry transferred to nitrocellulose membranes by a Trans-Blot Turbo Transfer System (Bio-Rad, Hercules, CA, USA). The membranes were blocked with either 5% bovine serum albumin in PBS or 5% (w/v) non-fat dried milk in TTBS at room temperature for 1 h and incubated overnight at 4 °C with specific anti-IL-1β (R&D Systems, Minneapolis, MN, USA), anti-phospho-NF-κB p65 (Ser536) (Cell Signaling, Danvers, MA, USA), anti-NF-κB p65 (Cell Signaling, Danvers, MA, USA), anti-phospho-mTOR (Ser2448) (Cell Signaling, Danvers, MA, USA) and anti-mTOR (Cell Signaling, Danvers, MA, USA) (concentration 1:1000) primary antibodies. Incubation with HRP-conjugated secondary antibodies (GE Healthcare, Chicago, IL, USA) was performed at room temperature for 1 h (dilution 1:10 000). Protein bands were developed by applying Clarity™ Western ECL Substrate (Bio-Rad, Hercules, CA, USA) for 5 min. Precision Plus Protein Kaleidoscope Standards (Bio-Rad, Hercules, CA, USA) were used for molecular weight determinations. Protein bands were visualized and imaged by using an ImageQuant LAS 500 scanner (GE Healthcare, Chicago, IL, USA). After transfer, the blots were stained with Ponceau S solution (Sigma‒Aldrich, St. Louis, MO, USA) and developed for total protein level normalization. Data were analyzed using the Image Studio^TM^ Lite software (LI-COR Biosciences, Lincoln, NE, USA).

### Statistical analysis

Data are expressed as the mean ± SEM. The equality of variance and normality of the data were checked by the Brown-Forsythe test and Shapiro-Wilk test, respectively. Based on the results, statistical analysis was performed using either t test (two groups), ordinary one-way ANOVA, Brown-Forsythe and Welch ANOVA or Kruskal-Wallis tests with correction for multiple comparisons by controlling the False Discovery Rate (two-stage linear step-up procedure of Benjamini, Krieger and Yekutieli) (Graphpad Prism 9.3.1, San Diego, CA, USA). Values of p < 0.05 (or q-values for proteomics experiments) were considered statistically significant.

## Results

### AP39 reduced weight and fatty liver in HFD-fed mice

In our experiments, we administered AP39 subcutaneously to C57BL/6J mice on a normal chow diet (control) and a high fat diet (HFD) three days a week in a dose of 100 nM per kg of body weight per day for 12 weeks (Fig. 1A). The mean weight of mice did not differ between control mice and control + AP39 mice at the end of experiment (Fig. 1C) and over 12 weeks of experiment duration (Fig. 1D). However, mice on an HFD significantly gained weight over 12 weeks of experiment duration (41.51 ± 1.51 g in HFD vs. 30.39 ± 0.51 g in control at the end of experiment; p<0.0001) (Fig. 1C and D) and AP39 administration reduced weight in HFD group starting from 6 week of treatment (35.98 ± 1.24 g in HFD + AP39 vs. 41.51 ± 1.51 g in HFD at the end of experiment; p<0.01) (Fig. 1C and D). Importantly, AP39 treatment did not influence the food consumption rate over 12 weeks of experiment (Fig. 1E). The action of AP39 as a donor of H_2_S was confirmed by the measurement of H_2_S level using methylene blue assay. The level of kit-detectable H_2_S was significantly higher in the liver (0.097 ± 0.009 μmol/gprot in control + AP39 vs. 0.059 ± 0.009 μmol/gprot in control; p<0.05) (Fig. 1F) of AP39-treated mice compared to control mice. At physiological pH, H_2_S (pKa 7.04) dissociates to the hydrosulphide anion (HS^−^) and the sulphide anion (S^2−^). Therefore, the term kit-detectable H_2_S refers to the sum of these species (H_2_S, HS^−^ and S^2−^) present at physiological pH ^29^.

**Figure 1.**
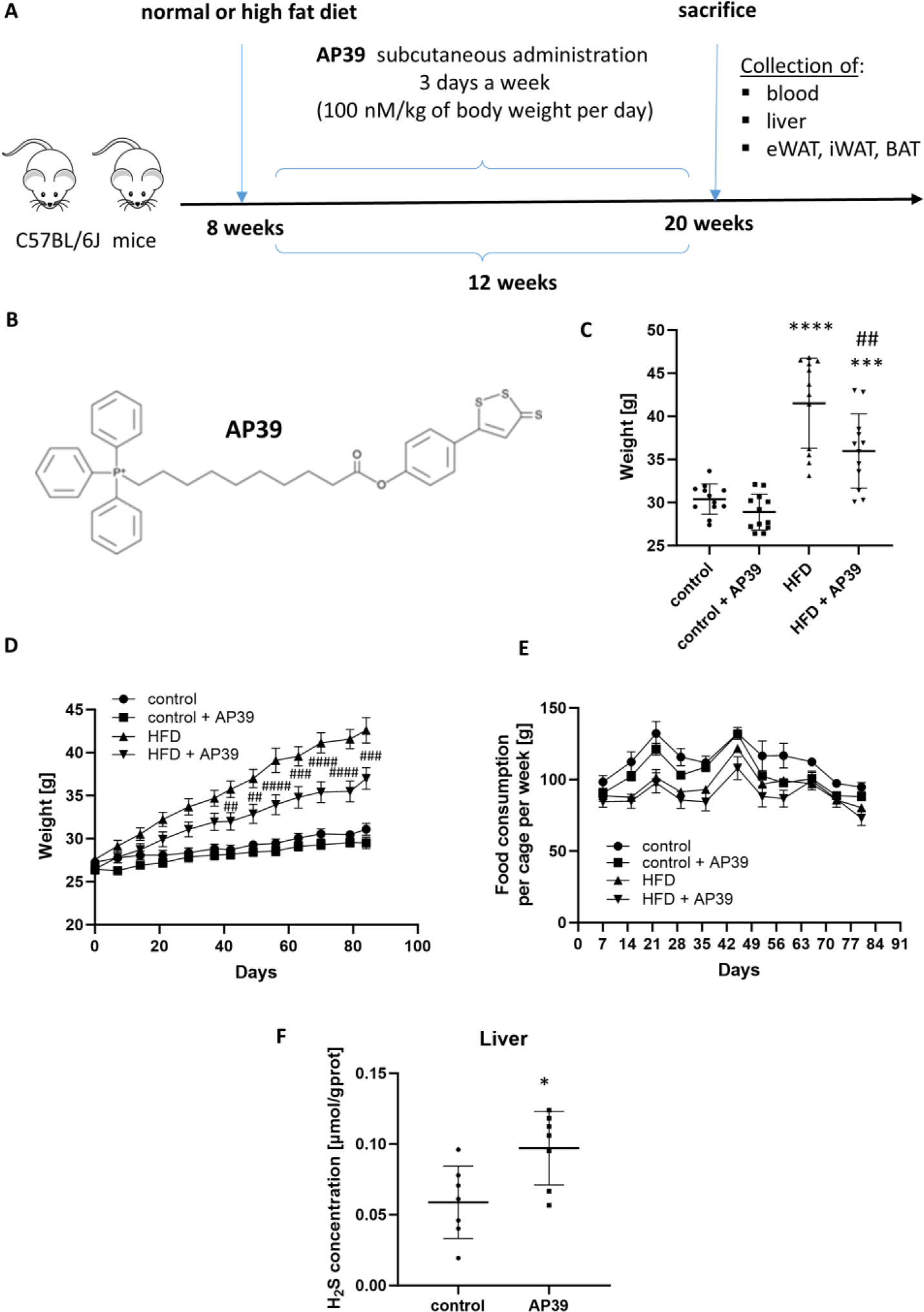
AP39 decreases weight of C57BL/6J mice fed a high fat diet. The experimental design of the study **(A)**. The chemical structure of AP39 **(B)**. Weight of control, control + AP39, HFD and HFD + AP39 groups at the end of experiment **(C)**. Weight of control, control + AP39, HFD and HFD + AP39 groups over 16 weeks of high-fat diet and drug administration **(D)**. Food consumption rate of control, control + AP39, HFD and HFD + AP39 groups over 16 weeks of diet and drug administration **(E)**. Concentration of H_2_S in the liver of control and AP39-treated mice **(F)**. Mean ± SEM; *p<0.05 compared to control mice; n=7 or 12.

Next, we assessed the impact of AP39 on the development of hepatic steatosis in the liver of C57BL/6J mice on HFD. The morphology of the liver was visually different between control and HFD-fed mice (Fig. 2A); however, the mean weight of the liver did not differ between groups (Fig. 2B). As shown by H&E staining, the cytoplasm of hepatocytes had a granular structure with signs of both macrovesicular and microvesicular steatosis of about 23% and 11% of hepatocytes, respectively (Fig. 2D, E and F). AP39 administration significantly reduced it to about 5% and 1% of hepatocytes, respectively (p<0.05) (Fig. 2D, E and F). The lobular structure of the liver was preserved with no inflammatory or necrotic changes. Treatment with AP39 also resulted in the significant decrease of triglycerides in the liver (0.20 ± 0.02 μg/mL in HFD + AP39 vs. 0.28 ± 0.01 μg/mL in HFD; p<0.0001) (Fig. 2C) and plasma (Table 1) of HFD-fed mice. AP39 administration did not impact plasma levels of total cholesterol, HDL, and LDL (Table 1) as well as plasma levels of ALT and AST enzymes, markers of liver damage (Fig. 2G and H). However, it increased plasma GSH/GSSG ratio in HFD-fed mice (3.83 ± 0.14 in HFD + AP39 vs. 3.24 ± 0.10 in HFD; p<0.05) (Fig. 2I), an indicator of oxidative stress and cellular health, with a higher ratio meaning less oxidative stress.

**Figure 2.**
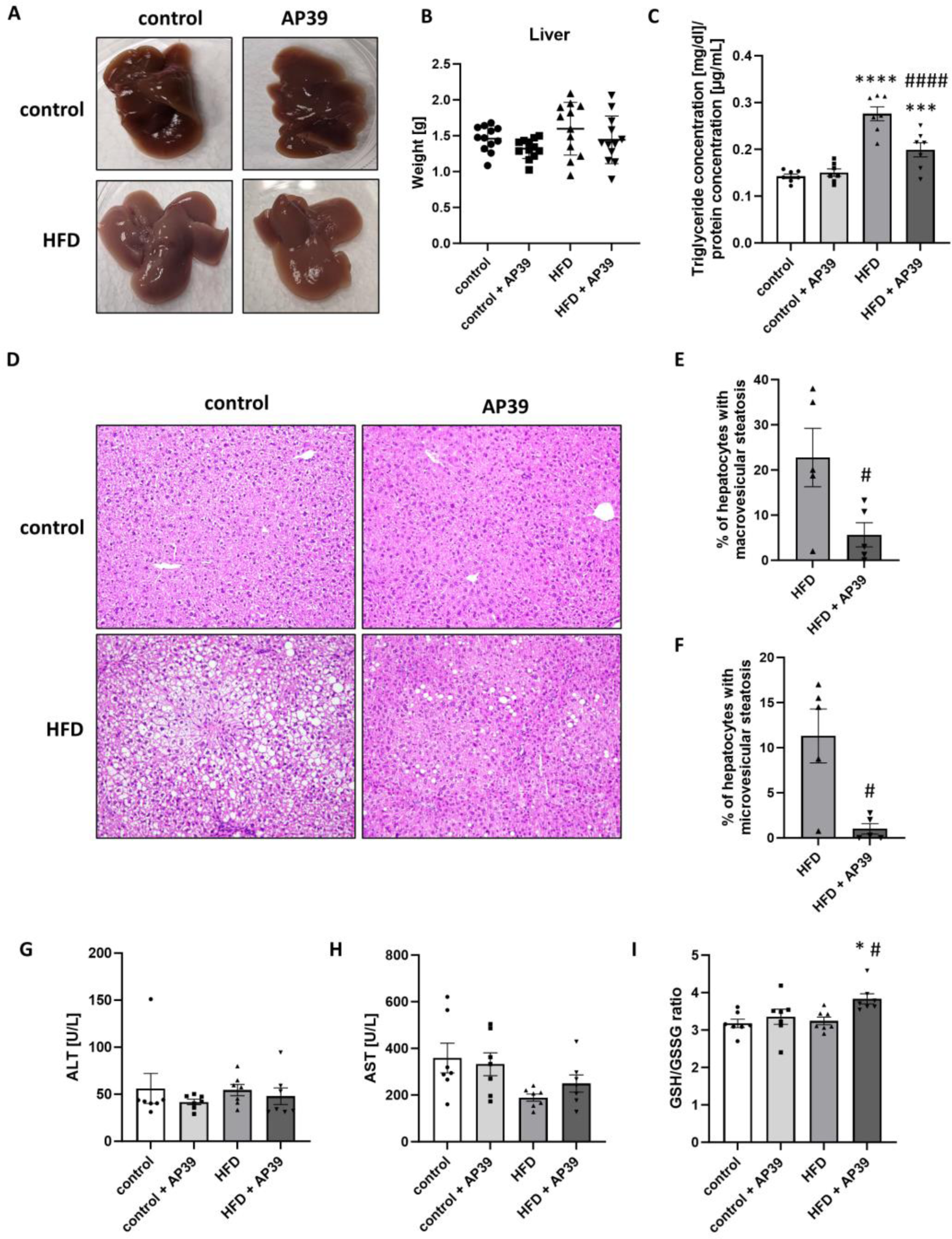
AP39 reduces fatty liver in HFD-fed mice. Representative images of the liver **(A)**, liver weight **(B)** and triglyceride content in the liver **(C)** of control, control + AP39, HFD and HFD + AP39 groups. Representative hematoxilin and eosin staining of the liver of control, control + AP39, HFD and HFD + AP39 groups **(D)**. Percent of hepatocytes with macrovesicular **(E)** and microvesicular steatosis **(F)** in mice fed an HFD and treated with AP39. Plasma levels of ALT **(G)**, AST (**H)** and plasma GSH/GSSG ratio **(I)** in control, control + AP39, HFD and HFD + AP39 groups. Mean ± SEM; *p<0.05, ***p<0.001, ****p<0.0001 compared to control group; #p<0.05, ####p<0.0001 compared to HFD group; n=5, 7 or 12.

**Table 1.** Plasma levels of total cholesterol, high-density lipoproteins (HDL), low-density lipoproteins (LDL), and triglycerides (TG) in control, control + AP39, HFD and HFD + AP39 groups, presented as mean ± SEM; n = 7 per group.

| Group | Total Cholesterol<br>[mmol/L] | HDL [mmol/L] | LDL [mmol/L] | TG [mmol/L] |
| --- | --- | --- | --- | --- |
| control | 2.31 ± 0.13 | 2.16 ± 0.12 | 0.31 ± 0.03 | 0.87 ± 0.05 |
| control +AP39 | 2.17 ± 0.20 | 1.94 ± 0.19 | 0.37 ± 0.04 | 0.70 ± 0.03** |
| HFD | 4.15 ± 0.51*** | 3.55 ± 0.45** | 0.69 ± 0.12* | 1.08 ± 0.07** |
| HFD + AP39 | 3.92 ± 0.32*** | 3.27 ± 0.28** | 0.69 ± 0.09* | 0.87 ± 0.04## |

\* $p$ <0.05, \*\* $p$ <0.01, \*\*\* $p$ <0.001 compared to control group; ## $p$ <0.01 compared to HFD group

### Proteomic analysis of the liver revealed downregulated pathways related to biosynthesis of unsaturated fatty acids, lipoprotein assembly and PPAR signaling upon AP39 administration in HFD-fed mice

To deeply investigate the differences in protein expression upon AP39 administration in the liver of HFD-fed mice, we conducted a quantitative proteomics analysis. LC‒MS/MS measurements operated in DIA mode and analyzed in Spectronaut software with directDIA^TM^ approach resulted in the identification and quantification of 6519 protein groups across all biological conditions. A summary of the quality control for the LC‒MS/MS runs is shown in Supplemental Figure 1. The median protein group CV was approximately 12% (Supplemental Figure 1D), which allowed for the calculation of a significant cutoff equal to a fold change of 1.25 (statistical power greater than 80%). Taken together, we found a total of 1689 proteins altered in the liver of HFD-fed mice compared to control group (Fig. 3A); 759 proteins altered between control + AP39 group and control group (Fig. 3B); and finally, 767 proteins changed in HFD + AP39 group compared to HFD group (Fig. 3C). Principal component analysis (PCA) showed clear data clustering between different biological groups (Supplemental Figure 1F). The detailed list of differentially expressed proteins and their fold changes across all biological conditions are presented in Supplemental Table 1.

**Figure 3.**
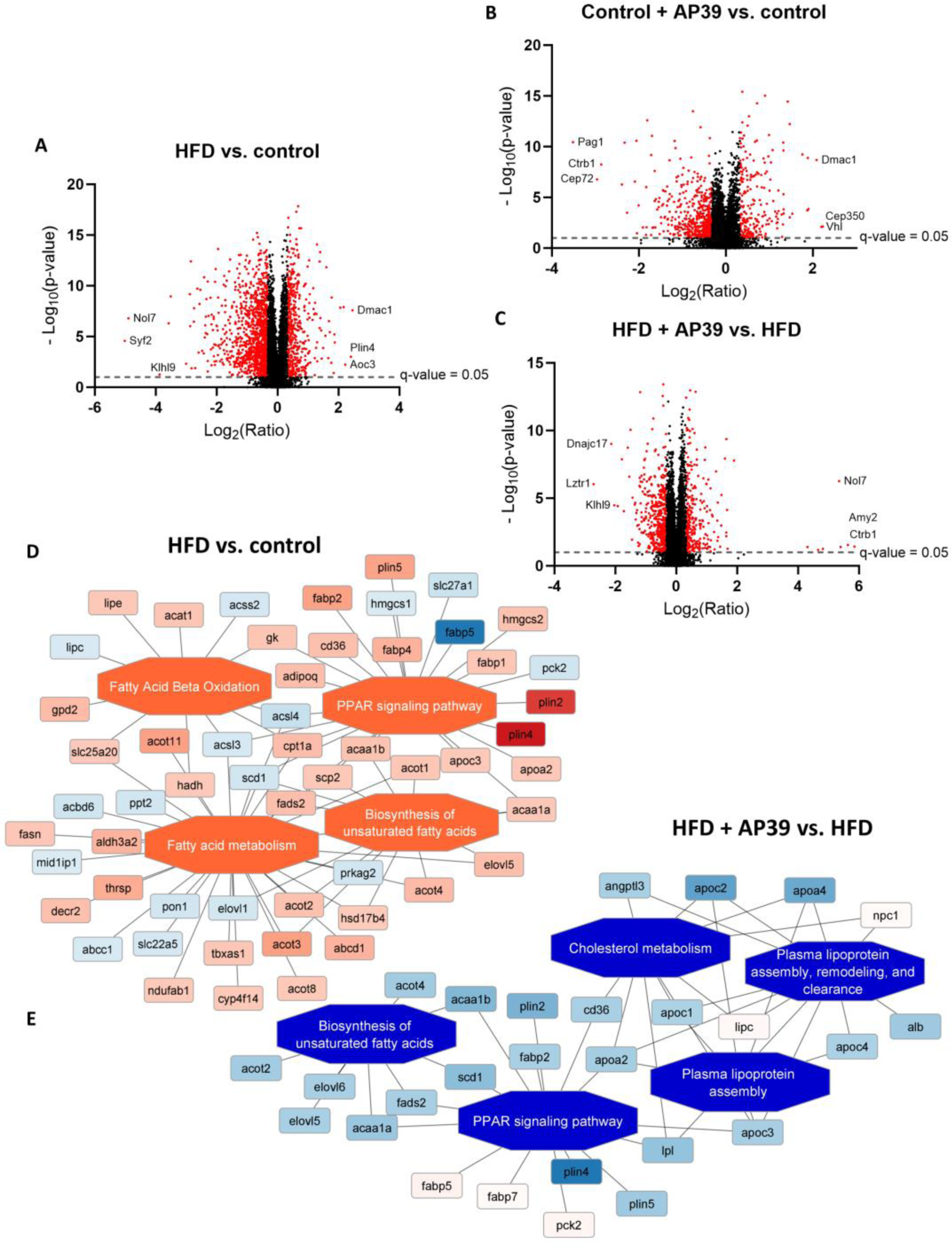
Proteomic analysis of the liver reveals downregulated pathways related to biosynthesis of unsaturated fatty acids, lipoprotein assembly and PPAR signaling in HFD-fed mice upon treatment with AP39. Volcano plot of differentially expressed proteins showing the log2 ratio of protein expression versus - log10 p value in HFD group compared to control **(A),** control + AP39 group compared to control **(B)** and HFD + AP39 group compared to HFD group **(C)**. The most upregulated and downregulated proteins are shown. Enriched functional network in HFD group compared to control **(D)** and HFD +AP39 group compared to HFD group **(E)**, as generated by PINE software **(B)**. Activated pathways are shown as orange central nodes, and inhibited pathways are shown as blue central nodes along with red (upregulated) or blue (downregulated) protein nodes. q-value<0.05; n=7.

To identify the functional pathways and biological networks that are altered in HFD-fed mice upon AP39 administration, we performed enriched pathway analysis based on protein expression using PINE software. Enriched functional networks indicated that pathways related to fatty acid metabolism, biosynthesis of unsaturated fatty acid, and peroxisome proliferator-activated receptor (PPAR) signaling were upregulated in the liver of HFD-fed mice compared to control group, shown as orange central nodes (Fig. 3D). On the other hand, pathways involved in biosynthesis of unsaturated fatty acids, cholesterol metabolism, PPAR signaling, and plasma lipoprotein assembly, shown as blue central nodes, were downregulated upon AP39 treatment in the liver of HFD-fed mice (Fig. 3E), which pointed to the beneficial action of AP39 in reversing altered pathways due to HFD feeding.

Importantly, our results indicated that AP39 administration resulted in the favorable changes of several proteins involved in the development of hepatic steatosis. Expressions of proteins engaged in *de novo* lipogenesis, such as sterol regulatory element-binding protein (SREBP-1; fold change (FC) = −2.01), stearoyl-CoA desaturase 1 (SCD1; FC = −1.63), mid1-interacting protein 1 (MIG12; FC = −2.25), and c-Jun N-terminal kinase 1 (JNK1; FC = −1.34) were decreased in the liver of AP39-treated HFD-fed mice. In addition, proteins related to insulin signaling pathway that are upstream in mTOR pathway, such as insulin receptor substrate 1 (IRS-1; FC = −1.86), insulin-like growth factor I (IGF-1; FC = −2.09) and insulin-like growth factor-binding protein 2 (IGFBP2; FC = 2.48) were differentially expressed because of AP39 treatment. Moreover, upon AP39 administration different perilipins (PLIN), structural components of lipid droplets (PLIN2, 4 and 5; FC = −1.45 to −2.74), were downregulated in the liver of HFD group. Interestingly, treatment with AP39 resulted in the altered expression of proteins regulating inflammation, particularly NF-κB signaling pathway, in the liver of HFD-fed mice, such as mitogen-activated protein kinase (MAPK) kinase kinase 2 (MEKK2; FC = −1.63), I-kappa-B kinase alpha (IKK-α; FC = −1.28), serum amyloid A-1 protein (SAA1; FC = −3.30), leukocyte cell-derived chemotaxin-2 (LECT2; FC = −1.60), leukocyte elastase inhibitor A (SERPINB1; FC = 2.56), and TRAF2-binding protein (T2BP; FC-1.53) (Supplemental Table 1).

On the other hand, we observed increased expression of some structural proteins that form extracellular matrix, including different types of collagen (type I, IV, V, VI, XIV and XVIII; FC = 1.26 to 2.80), different subunits of laminin (FC = 1.66 to 2.58) and nidogens (nidogen-1 and −2; FC = 1.76 and 1.48) upon AP39 administration in the liver of HFD-fed mice (Supplemental Table 1).

### AP39 decreased de novo lipogenesis and inflammation in the liver of HFD-fed mice

Noteworthy, our data pointed out to the downregulation of sterol regulatory element-binding transcription factor 1 (*Srebf1*), a crucial transcription factor responsible for the induction of *de novo* lipogenesis in the liver, upon AP39 administration in HFD-fed mice both at gene (FC = 1.40 ± 0.11 in HFD + AP39 vs. 2.36 ± 0.38 in HFD; p<0.05) (Fig. 4A) and protein level (FC = 0.67 ± 0.15 in HFD + AP39 vs. 1.34 ± 0.22 in HFD; p<0.001) (Fig. 4B). Simultaneously, we observed decreased mRNA expression of fatty acid synthase (*Fasn*), a pivotal enzyme participated in fatty acid synthesis, in the liver of AP39-treated HFD-fed mice (FC = 0.73 ± 0.17 in HFD + AP39 vs. 1.58 ± 0.33 in HFD; p<0.05) (Fig. 4C). Moreover, the expression of SCD-1 protein, which is a rate-liming enzyme in biosynthesis of unsaturated fatty acids, was decreased in the liver of AP39-treated HFD-fed mice (FC = 0.38 ± 0.13 in HFD + AP39 vs. 0.62 ± 0.12 in HFD; p<0.001) (Fig. 4D). Importantly, we demonstrated that AP39 inhibited mTOR signaling pathway, which was assessed by decreased phosphorylation of mTOR (Ser2448) normalized to total mTOR (0.098 ± 0.024 in HFD + AP39 vs. 0.186 ± 0.021 in HFD; p<0.05) (Fig. 4M, N, O; Supplemental Figure 2).

**Figure 4.**
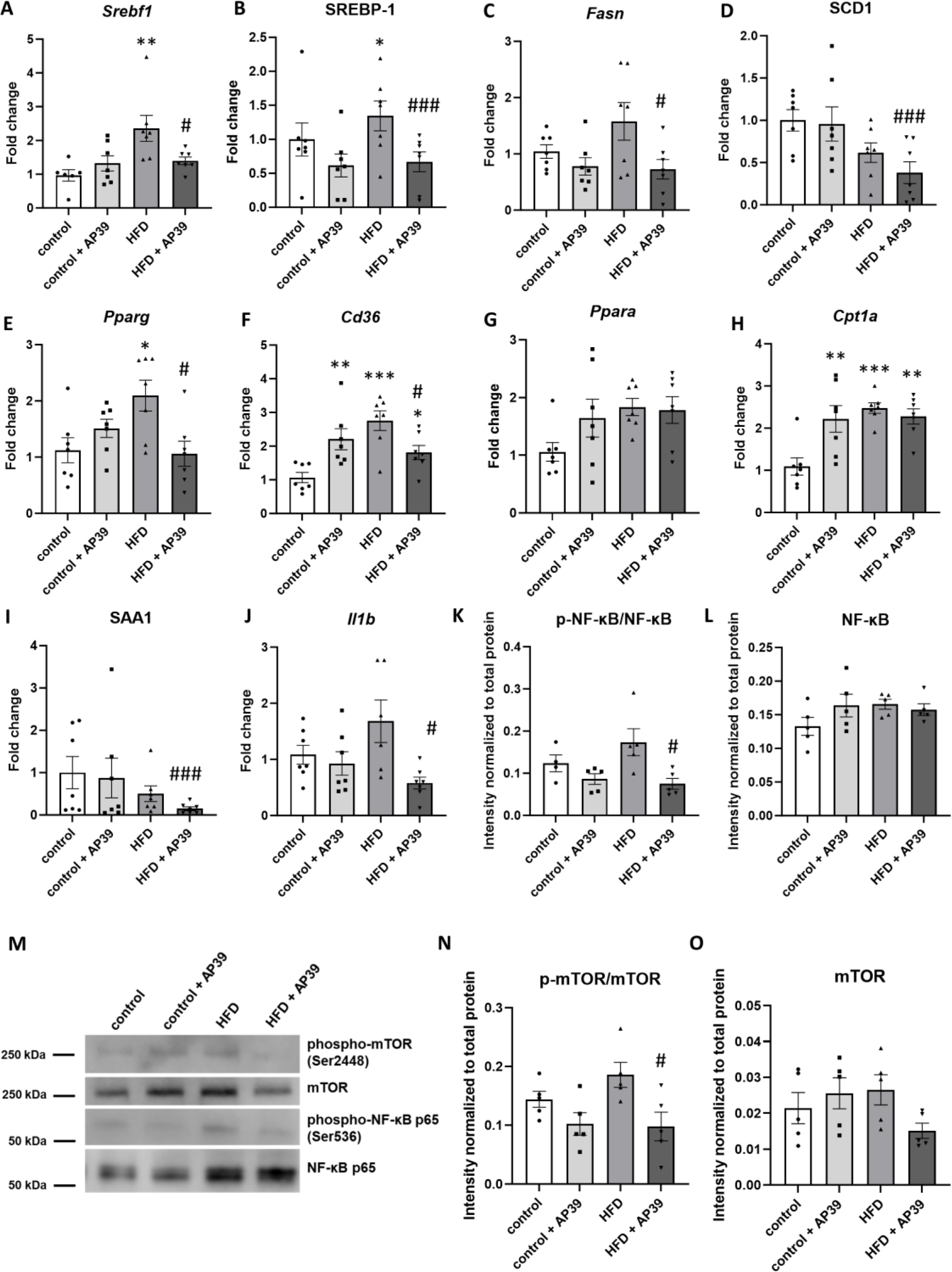
AP39 decreases *de novo* lipogenesis and inflammation in the liver of HFD-fed mice through downregulation of SREBP-1 transcription factor, SCD1 protein expression and NF-κB signaling pathway. mRNA expression of transcription factor *Srebf1* (A) and its protein expression (SREBP-1) **(B)** in the liver of control, control + AP39, HFD and HFD + AP39 groups. *Fasn* mRNA expression **(C)**, SCD1 protein expression **(D)** as well as mRNA expression of *Pparg* (E), *Cd36* (F), *Ppara* (G), *Cpt1a* (H) in the liver of control and HFD mice treated with AP39. Protein expression of SAA1 **(I)** and mRNA expression of *Il1b* (J) in the liver of control, control + AP39, HFD and HFD + AP39 groups. Western blot of phospho-mTOR (Ser2448), total mTOR, phospho-NF-κB p65 (Ser536) and total NF-κB p65 **(M)** as well as quantitative analysis of phospho-NF-κB p65 (Ser536) **(K)**, NF-κB p65 **(L)**, phospho-mTOR (Ser2448) **(N)** and mTOR **(O)** expressions in the liver of control and HFD mice treated with AP39. Mean ± SEM; *p<0.05, **p<0.01, ***p<0.001, ****p<0.0001 compared to control group; #p<0.05, ##p<0.01, ###p<0.001, ####p<0.0001 compared to HFD group; n=5-7.

In addition, our data showed downregulated mRNA expression of a transcription factor regulating fatty acid storage and glucose metabolism -peroxisome proliferator-activated receptor gamma (*Pparg*) (FC = 1.06 ± 0.22 in HFD + AP39 vs. 2.09 ± 0.28 in HFD; p<0.05) (Fig. 4E) and protein engaged in the import of fatty acids inside cells - cluster of differentiation 36 (*Cd36*) (FC = 1.81 ± 0.21 in HFD + AP39 vs. 2.76 ± 0.29 in HFD; p<0.05) (Fig. 4F) in the liver of HFD-fed mice upon AP39 treatment. Interestingly, AP39 administration solely in control mice resulted in increased mRNA expression of *Cd36* and carnitine palmitoyltransferase 1A (*Cpt1a*) that transports long chain fatty acids to mitochondria for fatty acid beta-oxidation (Fig. 4F and H). However, mRNA expression of a transcription factor responsible for fatty acid beta-oxidation - peroxisome proliferator-activated receptor alpha (*Ppara*) was not altered upon AP39 treatment (Fig. 4G).

Our results also showed that AP39 reduced inflammation in the liver of HFD-fed mice. mRNA expression of interleukin-1 beta (*Il1b*) (FC = 0.58 ± 0.11 in HFD + AP39 vs. 1.68 ± 0.38 in HFD; p<0.05) (Fig. 4J) as well as the activation of NF-κB signaling pathway, which was assessed by the measurement of phospho-NF-κB p65 (Ser536) normalized to total NF-κB p65 (0.075 ± 0.013 in HFD + AP39 vs. 0.174 ± 0.032 in HFD; p<0.05) (Fig. 4M, K, L; Supplemental Figure 2), were decreased upon AP39 administration in HFD group. Furthermore, protein expression of SAA1, which worsens hepatic steatosis via activating NF-κB signaling pathway, was downregulated in AP39-treated HFD-fed mice (FC = 0.15 ± 0.05 in HFD + AP39 vs. 0.50 ± 0.19 in HFD; p<0.001) (Fig. 4I).

### AP39 reduced obesity in HFD-fed mice

Apart from mitigating hepatic steatosis, AP39 also decreased obesity in HFD-fed mice. We clearly observed hypertrophy of white adipocytes in eWAT and iWAT of HFD-fed mice, the adipocyte diameters were significantly larger in HFD-fed mice than control mice. We found that AP39 administration reduced the diameters of adipocytes in eWAT (101.4 ± 1.62 μm in HFD + AP39 vs. 135.4 ± 4.39 μm in HFD; p<0.0001)(Fig. 5A and C), weight of eWAT (1.72 ± 0.13 g in HFD + AP39 vs. 2.32 ± 0.19 g in HFD; p<0.01) (Fig. 5B) as well as the number of foci of inflammation (0.6 ± 0.25 in HFD + AP39 vs. 5.2 ± 1.43 in HFD; p<0.01)(Fig. 5D) in eWAT of HFD-fed mice. Moreover, AP39 treatment reduced weight of iWAT (91.38 ± 6.04 g in HFD + AP39 vs. 98.50 ± 2.69 g in HFD; p<0.05) (Fig. 5F) but did not influence the diameters of adipocytes in iWAT of HFD group (Fig. 5E and G). Similarly, we did not see any changes in weight (Fig. 5I) and morphology of BAT (Fig. 5H and J) upon AP39 administration in HFD group. In addition, we did not observe any foci of inflammation in iWAT and BAT of HFD-fed mice.

**Figure 5.**
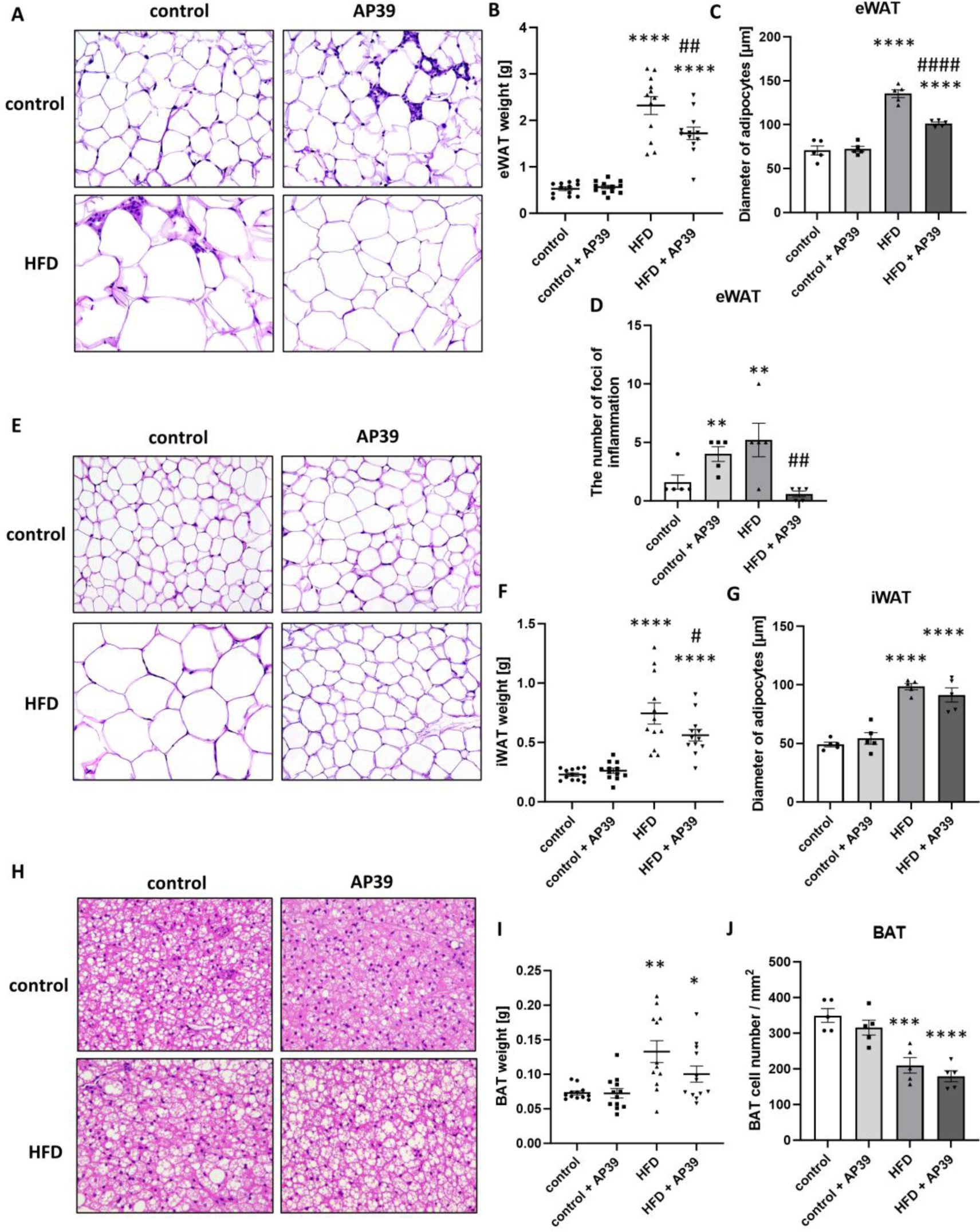
AP39 reduces obesity in HFD-fed mice. Representative hematoxilin and eosin (H&E) staining of eWAT **(A)**, eWAT weight **(B)** diameter of eWAT lipid droplets **(C)** and the number of foci of inflammation **(D)** in eWAT of control, control + AP39, HFD and HFD + AP39 groups. Representative H&E staining of iWAT **(E)**, iWAT weight **(F)** diameter of iWAT lipid droplets **(G)** in control and HFD mice treated with AP39. Representative H&E staining of BAT **(H)**, BAT weight **(I)** and BAT cell number / mm^2^ **(J)** in control, control + AP39, HFD and HFD + AP39 groups. Mean ± SEM; *p<0.05, **p<0.01, ***p<0.001, ****p<0.0001 compared to control group; #p<0.05, ##p<0.01, ####p<0.0001 compared to HFD group; n=5 or 12.

The alleviation of obesity caused by AP39 in HFD-fed mice was not related to the improvement of insulin resistance in these mice. As shown by glucose tolerance test (GTT) (Fig. 6A), there was no difference between HFD group and HFD + AP39 group, which was measured as the area under the curve (AUC) (Fig. 6B). Moreover, fasting blood glucose levels (Fig. 6C), fasting plasma insulin levels (Fig. 6D) and HOMA-IR insulin resistance index (Fig. 6E) were not altered upon AP39 administration in HFD-fed mice. However, as expected, all these parameters were significantly higher in HFD-fed mice in comparison to control group, which indicated increased insulin resistance upon a high fat diet.

**Figure 6.**
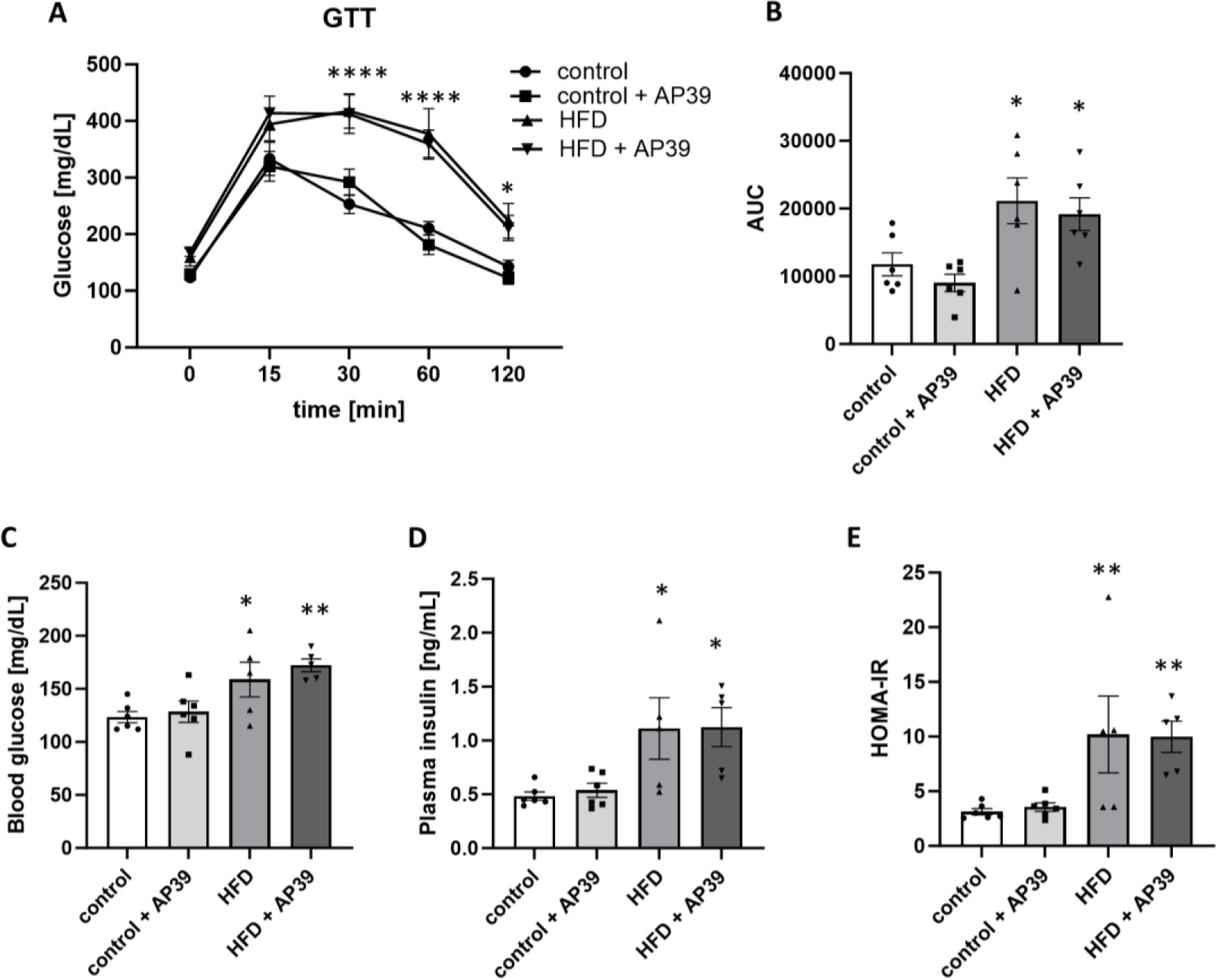
AP39 does not influence insulin resistance in HFD-fed mice. Glucose tolerance test (GTT) **(A)** and the area under the curve (AUC) **(B)** of control and HFD mice treated with AP39 for 12 weeks. Fasting blood glucose level **(C)**, fasting plasma insulin level **(D)** and HOMA-IR insulin resistance index **(E)** of control, control + AP39, HFD and HFD + AP39 groups. Mean ± SEM; *p<0.05, **p<0.01, ****p<0.0001 compared to control group; n=5-6.

### AP39 decreased inflammation in adipose tissue of HFD-fed mice

To further explore the impact of AP39 on the reduction of obesity, particularly observed in eWAT, we measured expression of proinflammatory genes as well as genes regulating hydrolysis and synthesis of triglycerides in eWAT. mRNA expression of *Il1b* (FC = 1.14 ± 0.24 in HFD + AP39 vs. 4.10 ± 0.96 in HFD; p<0.001) (Fig. 7A), *Il6* (FC = 3.35 ± 0.70 in HFD + AP39 vs. 7.04 ± 1.56 in HFD; p<0.05) (Fig. 7B), *Tnf* (FC = 3.10 ± 0.70 in HFD + AP39 vs. 6.94 ± 1.10 in HFD; p<0.001) (Fig. 7C) and *Mcp1* (FC = 8.25 ± 2.53 in HFD + AP39 vs. 20.00 ± 5.27 in HFD; p<0.05) (Fig. 7D) was significantly decreased upon AP39 administration in HFD-fed mice. Moreover, protein expression of pro-IL-1β (0.067 ± 0.002 in HFD + AP39 vs. 0.99 ± 0.003 in HFD; p<0.0001) (Fig. 7L and M; Supplemental Figure 3) was also diminished in eWAT of AP39-treated HFD-fed mice. Intriguingly, we observed that AP39 treatment in control mice on normal chow diet upregulated mRNA expression of some proinflammatory markers including *Il1b* (FC = 2.03 ± 0.41 in control + AP39 vs. 1.06 ± 0.15 in control; p<0.05) (Fig. 7A) and *Mcp1* (FC = 2.19 ± 0.30 in control + AP39 vs. 1.03 ± 0.10 in control; p<0.05) (Fig. 7D) in eWAT. Concomitantly, the number of foci of inflammation was increased in eWAT of control mice upon AP39 administration (4.00 ± 0.63 in control + AP39 vs. 1.60 ± 0.60 in control; p<0.01) (Fig. 5D).

**Figure 7.**
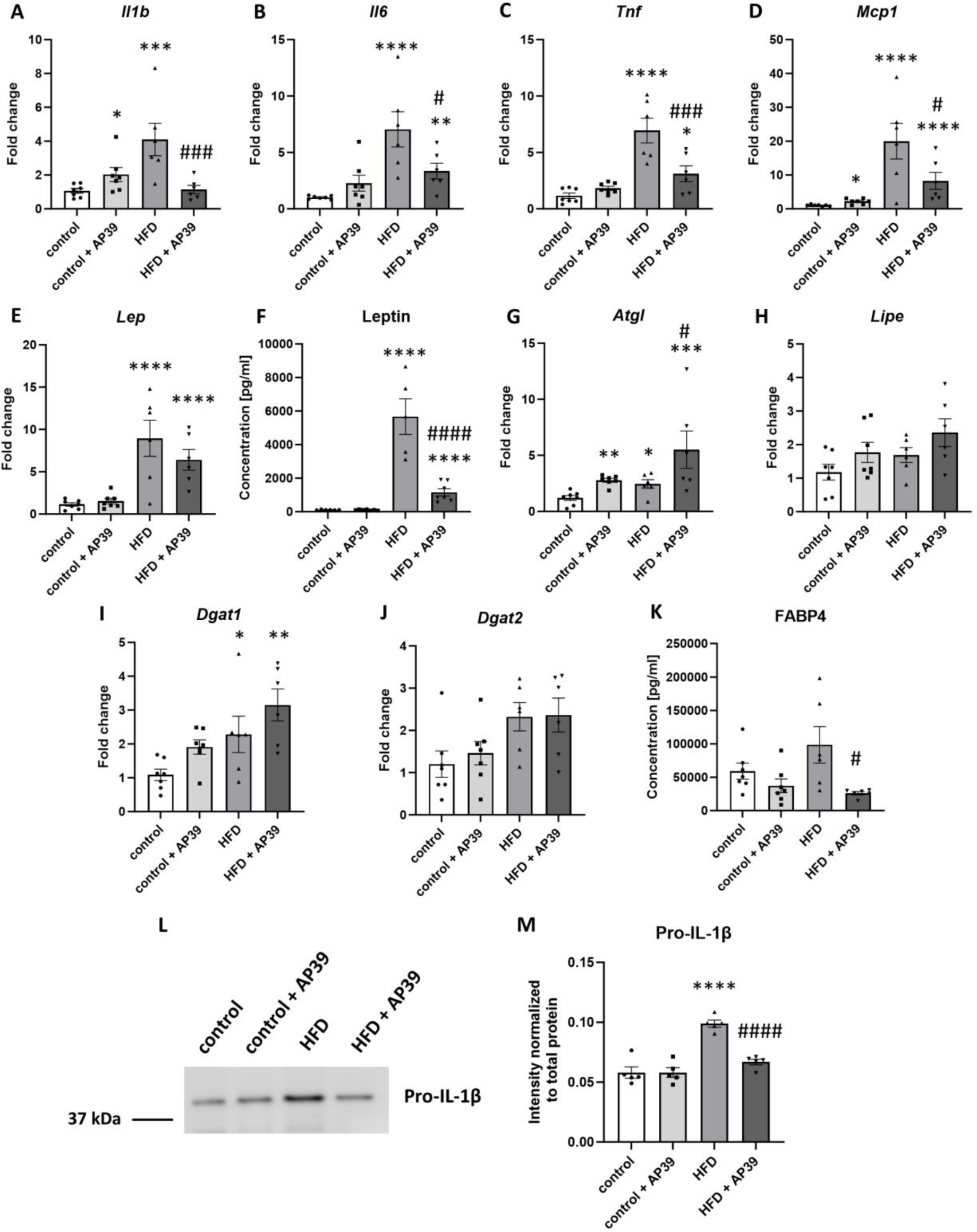
AP39 decreases inflammation and leptin level in HFD-fed mice. mRNA expression of inflammatory genes such as *Il1b* **(A)**, *Il6* **(B)**, *Tnf* **(C)** and *Mcp1* **(D)** in eWAT of control, control + AP39, HFD and HFD + AP39 groups. *Lep* mRNA expression in eWAT **(E)** as well as leptin plasma level **(F)** in control and HFD mice treated with AP39. mRNA expression of genes regulating hydrolysis and synthesis of triglycerides in adipocytes including *Atgl* **(G)**, *Lipe* **(H)**, *Dgat1* **(I)** and *Dgat2* **(J)** in eWAT of control, control + AP39, HFD and HFD + AP39 groups. FABP4 protein plasma level across all biological conditions **(K)**. Western blot of pro-IL-1β **(L)** as well as its quantitative analysis (**M**) in eWAT of control and HFD mice treated with AP39. Mean ± SEM; *p<0.05, **p<0.01, ***p<0.001, ****p<0.0001 compared to control group; #p<0.05, ###p<0.001, ####p<0.0001 compared to HFD group; n=5-7.

In line with our results on the reduction of obesity by AP39, our data also pointed to the decrease of leptin levels in the plasma of AP39-treated HFD-fed mice (1156 ± 216 pg/mL in HFD + AP39 vs. 5667 ± 1065 pg/mL in HFD; p<0.0001) (Fig. 7F) but not leptin mRNA expression (*Lep*) in eWAT (Fig. 7E). As expected, leptin expression was significantly increased in HFD-fed mice in comparison to control group both at mRNA and protein levels (Fig. 7 E and F). Furthermore, we showed that AP39 treatment upregulated mRNA expression of adipose triglyceride lipase (*Atgl*) (FC = 5.50 ± 1.67 in HFD + AP39 vs. 2.45 ± 0.38 in HFD; p<0.05) (Fig. 7G), an enzyme that hydrolyses triglycerides, in eWAT of HFD-fed mice. mRNA expression of another enzyme involved in lipolysis – hormone-sensitive lipase (*Lipe*) (Fig. 7H) as well as mRNA expression of enzymes engaged in the synthesis of triglycerides (diacylglycerol O-acyltransferase 1/2, *Dgat1*/*Dgat2*) (Fig. 7I and J) did not differ in eWAT of HFD-fed mice upon AP39 administration. Interestingly, AP39 treatment caused a decline in plasma protein level of fatty acid binding protein 4 (FABP4) (25934 ± 2406 pg/mL in HFD + AP39 vs. 98510 ± 27263 pg/mL in HFD; p<0.05) (Fig. 7K), which is a biomarker of obesity and metabolic syndrome.

## Discussion

Mounting evidence points to the important role of H_2_S in the physiological and pathological processes in the liver, including lipid metabolism and fibrosis. In the present study, we applied direct delivery of H_2_S to mitochondria using mitochondria-targeted H_2_S donor AP39 to evaluate the development of fatty liver and obesity in C57BL/6J mice fed an HFD. We demonstrated for the first time that AP39 treatment in HFD-fed mice **1)** reduced hepatic steatosis and obesity, **2)** decreased *de novo* lipogenesis in the liver by downregulating mTOR/SREBP-1/SCD1 signaling pathway, **3)** and decreased inflammation in the liver and eWAT by downregulating NF-κB signaling pathway.

Our results showed that AP39 administrated at a low dose (100 nM/kg/day) for 12 weeks significantly reduced weight of HFD-fed mice starting from 6 week of treatment without decreasing food consumption rate. This was accompanied by the reduction of macro-vesicular and micro-vesicular steatosis in the liver as well as triglyceride level in the plasma and liver of HFD-fed mice. However, AP39 treatment did not influence plasma levels of ALT and AST enzymes, markers of liver damage. On the other hand, it reduced oxidative stress measured as plasma GSH/GSSG ratio, which is an indicator of oxidative stress and cellular health. This is in line with other studies pointing to the importance of H_2_S signaling in the pathogenesis of fatty liver. Hepatic-specific CSE knockout was shown to promote the development of fatty liver in HFD-fed mice ^30^. In contrast, the deletion of 3-MST ameliorated hepatic steatosis in HFD-fed mice. 3-MST, both cytosolic and mitochondrial H_2_S-producting enzyme, could interact with CSE and inhibit its function, as a main source of H_2_S in the liver^31^. Moreover, it was observed that administration of exogenous H_2_S in the form of inorganic salts (NaSH) or slow-releasing H_2_S donor (GYY4137) reduced fatty liver by activating autophagy through AMPK/mTOR pathway ^32,33^, improving antioxidant potential ^34^ and decreasing hepatic endoplasmic reticulum stress ^35^. However, in the present study, we exploited the direct and controlled delivery of H_2_S to mitochondria using mitochondria-targeted H_2_S donor AP39 to investigate the development of fatty liver and obesity in HFD-fed mice.

The proteomics approach allowed us to discover downregulated pathways involved in biosynthesis of unsaturated fatty acids, cholesterol metabolism, PPAR signaling, and plasma lipoprotein assembly upon treatment with AP39 in the liver of HFD-fed mice. Particularly, AP39 administration inhibited *de novo* lipogenesis in the liver through the downregulation of mTOR/SREBP-1/SCD1 signaling pathway. We found that AP39 reduced the activation of mTOR by decreasing its phosphorylation on Ser2448. Furthermore, it downregulated SREBP-1 expression at both the gene and protein levels in the liver of AP39-treated HFD-fed mice. SREBP-1 is a crucial transcription factor responsible for regulating the expression of genes involved in *de novo* lipogenesis, such as FASN and SCD1 ^36^. SREBP-1 activation was shown to be important in the pathogenesis of fatty liver and its levels were increased in the liver of HFD-fed mice and patients with MAFLD ^37^. SREBP-1 expression was also upregulated in the liver of HFD-fed mice in our study. Noteworthy, it was demonstrated that exogenous H_2_S could enhance the degradation of SREBP-1 by ubiquitin-proteasome pathway ^38^. Consistently, our results pointed to downregulated expression of *Fasn* and SCD1 upon AP39 administration in the liver of HFD-fed mice. FASN is a major enzyme engaged in *de novo* lipogenesis that catalyzes the synthesis of palmitate from acetyl-CoA and malonyl-CoA, whereas SCD1 is a key rate-limiting enzyme in lipid metabolism by forming monounsaturated fatty acids (MUFAs) from saturated fatty acids (SFAs). SCD1 serves as a metabolic hub connecting pathways related to lipid metabolism, autophagy, and inflammation. Elevated levels of SCD1 were associated with increased risk of MAFLD, diabetes and obesity ^39^. Moreover, knockout of SCD1 in mice rendered them resistant to HFD-induced obesity ^40^ and prevented hepatic steatosis ^41^, which was related to suppressed lipogenesis and enhanced fatty acid oxidation. Apart from reduced expression of SREBP-1, we also found decreased expression of *Pparg* in the liver of AP39-treated HFD-fed mice. PPARγ is an another important transcription factor regulating lipogenesis (FASN expression), lipid uptake (upregulation of CD36 and FABP4) and formation of lipid droplets (upregulation of perilipins) in hepatocytes, thus promoting hepatic steatosis ^42^. Interestingly, our data showed reduced expression of *Cd36* and perilipins in the liver as well as decreased expression of FABP4, which is a biomarker of obesity and metabolic syndrome, in the plasma of HFD-fed mice upon AP39 administration.

Moreover, our proteomic data pointed out to the beneficial changes in expression of several proteins associated with mTOR-*de novo* lipogenesis pathway. We observed decreased expression of MIG12 and JNK1 in the liver of AP39-treated HFD-fed mice. MIG12 mediates stimulation of lipogenesis by LXR activation and its depletion leads to diminished expression of SREBP-1 and FASN ^43^. In turn, JNK1 promotes the progression of diet-induced MAFLD by elevating lipogenesis and decreasing fatty acid oxidation ^44^. Importantly, our proteomic results also revealed differentially expressed proteins related to insulin signaling pathway that are upstream in mTOR pathway. We found downregulated expression of IRS-1 and IGF-1 as well as upregulated expression of IGFBP2 in the liver of HFD-fed mice upon AP39 treatment. IRS-1 is an adaptor signaling protein that transmits signals from the insulin receptor to intracellular PI3K/AKT/mTOR pathway, whereas IGF-1 is a growth factor structurally and functionally related to insulin that binds to insulin receptor and activates PI3K/AKT/mTOR pathway ^45^. In addition, IGFBP2 serves as a transport protein for IGF-1 and mice with overexpressed IGFBP2 were resistant to the development of obesity ^46^. Altogether, our data pointed out to the inhibition of *de novo* lipogenesis in the liver of HFD-fed mice by AP39 through the downregulation of mTOR/SREBP-1/SCD1 signaling pathway.

Notably, our results also indicated that AP39 treatment decreased inflammation in the liver of HFD-fed mice, which was related to the downregulation of NF-κB signaling pathway. We showed that AP39 reduced activation of NF-κB pathway measured as decreased phosphorylation on Ser536 of NF-κB p65 as well as decreased *Il1b* expression in the liver. Interestingly, it was demonstrated that mTOR is an upstream regulator of NF-κB pathway ^47,48^ and in the present study we also observed reduced mTOR pathway upon AP39 administration. Furthermore, our MS data revealed diminished expression of several proteins involved in NF-κB pathway, such as MEKK2, IKK-α, SAA1 and T2BP. MEKK2 is an upstream serine/threonine protein kinase in NF-κB signaling pathway that directly phosphorylates and activates IκB kinases (IKK). IKK-α is a part of IκB kinase complex that phosphorylates IκBα protein, which leads to its degradation and subsequent activation of NF-κB pathway ^49^. In turn, SAA1, an acute-phase response protein, was reported to aggravate fatty liver through direct activation of TLR4-mediated NF-κB signaling pathway ^50^. SAA1 knockout was shown to reduce obesity and insulin resistance via inhibition of NF-κB pathway as well ^51^. Similarly, T2BP, which is a TRAF2 binding protein, was demonstrated to activate NF-κB signaling pathway ^52^. It is worth noting that all above proteins were downregulated upon AP39 administration in the liver of HFD-fed mice. Moreover, our proteomic results also pointed out to altered expression of two hepatokines, downregulated LECT2 and upregulated SERPINB1 in the liver of AP39-treated, HFD-fed mice. LECT2 level was shown to increase in the serum of patients with obesity and fatty liver ^53^. LECT2 links hepatic steatosis to inflammation via sensing the level of fat in the liver and shifting macrophages to proinflammatory M1 phenotype, thus contributing to the development of liver inflammation ^54^. SERPINB1 governs innate immune response, particularly negatively regulates IL-1β production ^55^. In addition, higher serum levels of SERPINB1 were associated with lower risk of type 2 diabetes mellitus^56^. Taking together, decreased expression of proteins engaged in NF-κB signaling pathway may contribute to the beneficial action of AP39 in the reduction of liver inflammation and steatosis.

Finally, we found that AP39 reduced obesity and inflammation in adipose tissue of HFD-fed mice. It was related to the reduction of eWAT and iWAT weights as well as adipocyte diameters and foci of inflammation in eWAT, but not iWAT and BAT. Intriguingly, AP39 administration did not influence insulin resistance measured by GTT, fasting blood glucose level, fasting plasma insulin level and HOMA-IR insulin resistance index. However, it decreased plasma level of leptin, a hormone that regulates a long-term energy balance and weight maintenance, in HFD-fed mice. Interestingly, it was shown that the reduction of leptin level is compulsory for weight loss, thus being a good strategy in the treatment of obesity ^57^. Moreover, we observed that AP39 administration downregulated the expression of markers related to inflammation, such as *Il1b*, *Il6*, *Tnf* and *Mcp1* at gene level and IL-1β at protein level in eWAT of HFD-fed mice. Similarly, it was reported that H_2_S donor NaHS reduced HFD-induced obesity and inflammation in mouse hypothalamus through inhibition of mTOR/NF-κB signaling pathway ^48^. Importantly, obesity is associated with low-grade chronic inflammation, thus targeting adipose tissue inflammation could have a beneficial impact in the treatment of obesity and obesity-related metabolic disorders ^58^. Our results also showed upregulated expression of *Atgl* upon AP39 treatment in eWAT of HFD-fed mice. However, the expression of other enzymes involved in lipolysis and synthesis of triglycerides did not differ upon AP39 administration. ATGL is an enzyme that catalyzes the hydrolysis of triacylglycerols to diacylglycerols, the first step of lipolysis ^59^. It is tempting to speculate that AP39 could ameliorate obesity in HFD-fed mice by decreasing inflammation and increasing ATGL expression in eWAT, which was reflected by reduced eWAT weight, inflammation, and adipocyte diameters.

It should be noted that AP39 treatment in control mice exerted some adverse proinflammatory effects regarding the increase in foci of inflammation and *Il1b* and *Mcp1* expressions in eWAT. This might be related to the fact that H_2_S exhibits beneficial effects only in low concentrations and this range of low concentrations could be different for healthy and diseased tissues, because in many pathological conditions the level of H_2_S is disturbed. Furthermore, our proteomic approach revealed upregulated expressions of some structural proteins that form extracellular matrix, including collagens, laminins and nidogens upon AP39 administration in HFD-fed mice. It is speculative whether these changes are a result of compensatory mechanism related to reduced inflammation in the liver upon AP39 treatment or are an indicator of developing fibrosis. Generally, it was shown that H_2_S has anti-fibrotic effects in the liver by reducing inflammation, oxidative stress and regulating lipid metabolism^60^. Nevertheless, one paper demonstrated upregulation of collagen type I upon AP39 administration in UVA-induced skin aging *in vitro* model ^61^. Further research is needed to elucidate the impact of mitochondria-targeted sulfide on the development of fibrosis.

Overall, our study indicated that AP39 reduced fatty liver, obesity, and inflammation in HFD-fed mice. This effect was mediated by inhibition of NF-κB signaling pathway and *de novo* lipogenesis by downregulating mTOR/SREBP-1/SCD1 signaling pathway (Figure 8). It is therefore tempting to speculate that mitochondria-targeted sulfide could provide potentially a novel therapeutic approach to the treatment/prevention of fatty liver and obesity.

**Figure 8.**
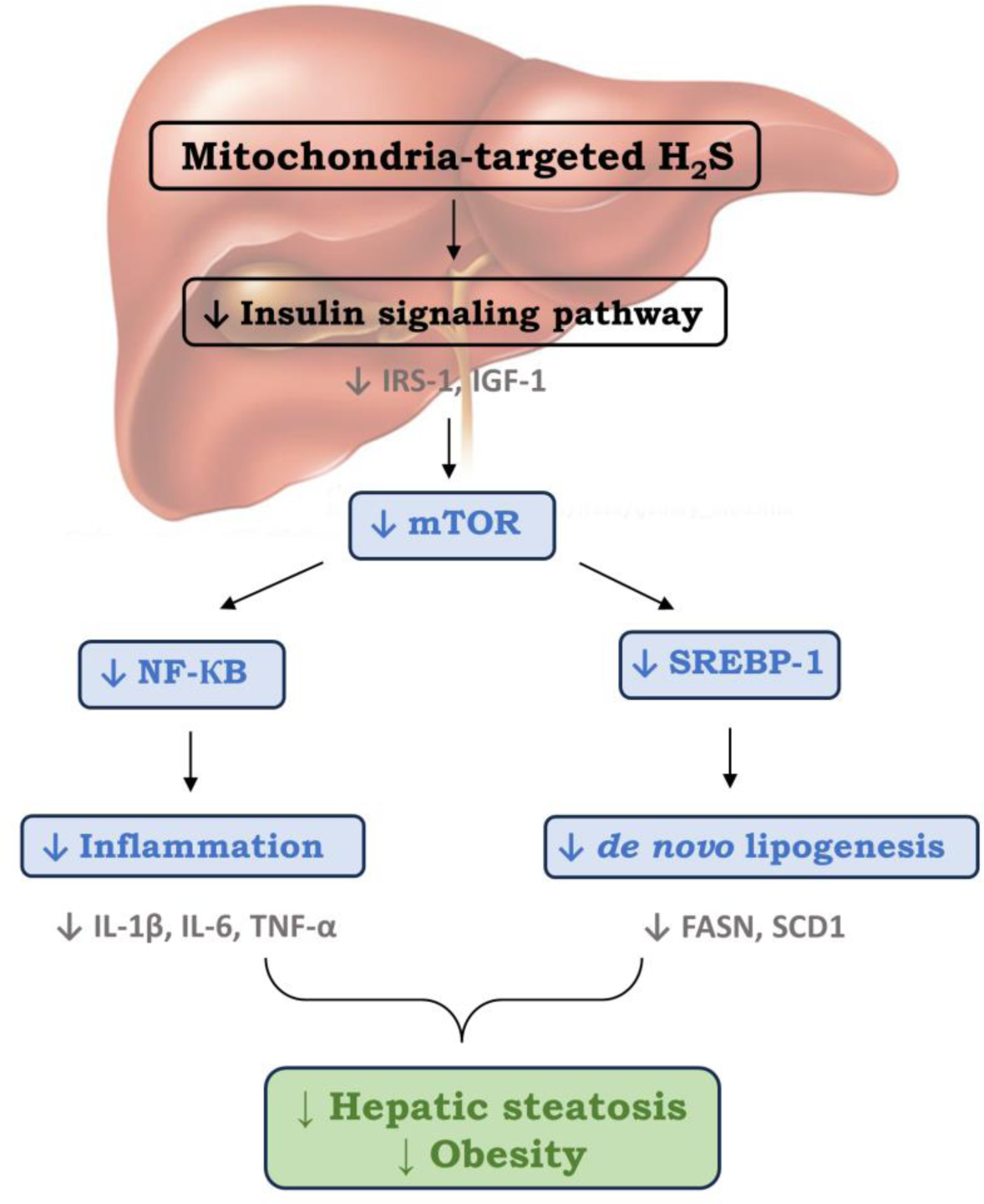
The summary of beneficial effects of mitochondria-targeted sulfide AP39 on the development of fatty liver and obesity.

## Conclusions

Our findings demonstrated that mitochondrial donor of H_2_S AP39 reduced fatty liver in HFD-fed mice, which was related to decreased level of TG in the liver and plasma as well as increased GSH/GSSG ratio in the plasma. It also inhibited *de novo* lipogenesis in the liver by downregulating mTOR/SREBP-1/SCD1 signaling pathway. Moreover, AP39 administration alleviated obesity in HFD-fed mice, which was reflected by reduced weight of mice and adipose tissue, decreased leptin level in the plasma and upregulated expression of ATGL, a lipolysis enzyme in eWAT. In addition, AP39 reduced inflammation in the liver and adipose tissue by downregulating mTOR/NF-κB signaling pathway. These results suggested that small molecule targeting of sulfide to mitochondria could be a novel therapeutic strategy in the treatment of obesity-related metabolic disorders. We do hope that knowledge regarding the mechanisms of action of mitochondria-targeted H_2_S can provide new insights for the pathogenesis of hepatic steatosis and obesity as well as facilitate the advancement of new therapeutic targets.

## Authors contribution

AS was responsible for the conception and design of the study. AS, AW, KC, KS, MUB, MS, BKC, GM, MW, RT, MW and KK were responsible for analyses of the samples. AS and RO were responsible for the interpretation of the data. AS drafted the article. All authors revised the paper critically for important intellectual content and gave final approval of the version to be published.

## Funding

This study was supported by the grant from National Science Centre (NCN): 2017/26/D/NZ4/00480. This research was carried out with the use of research infrastructure co-financed by the Smart Growth Operational Programme POIR 4.2 project no. POIR.04.02.00-00-D023/20.

## Conflict of interest

MW, RT, and MEW have intellectual property (patents awarded and pending) on slow-release sulfide-generating molecules and their therapeutic use. MW is CSO of MitoRx Therapeutics, Oxford, U.K, developing organelle-targeted molecules for clinical use.

## References

1. Chew NWS, Ng CH, Tan DJH, Kong G, Lin C, Chin YH, Lim WH, Huang DQ, Quek J, Fu CE, Xiao J, Syn N, Foo R, Khoo CM, Wang JW, Dimitriadis GK, Young DY, Siddiqui MS, Lam CSP, Wang Y, Figtree GA, Chan MY, Cummings DE, Noureddin M, Wong VWS, Ma RCW, Mantzoros CS, Sanyal A, Muthiah MD. The global burden of metabolic disease: Data from 2000 to 2019. Cell Metab. 2023;35(3):414–428.e3. doi:10.1016/j.cmet.2023.02.003

2. Xu H, Barnes GT, Yang Q, Tan G, Yang D, Chou CJ, Sole J, Nichols A, Ross JS, Tartaglia LA, Chen H. Chronic inflammation in fat plays a crucial role in the development of obesity-related insulin resistance. J Clin Invest. 2003;112(12):1821–1830. doi:10.1172/JCI19451

3. Kitade H, Chen G, Ni Y, Ota T. Nonalcoholic Fatty Liver Disease and Insulin Resistance: New Insights and Potential New Treatments. Nutrients. 2017;9(4):387. doi:10.3390/nu9040387

4. Eslam M, Sanyal AJ, George J, International Consensus Panel. MAFLD: A Consensus-Driven Proposed Nomenclature for Metabolic Associated Fatty Liver Disease. Gastroenterology. 2020;158(7):1999–2014.e1. doi:10.1053/j.gastro.2019.11.312

5. Cohen JC, Horton JD, Hobbs HH. Human fatty liver disease: old questions and new insights. Science. 2011;332(6037):1519-1523. doi:10.1126/science.1204265

6. Filipovic B, Marjanovic-Haljilji M, Mijac D, Lukic S, Kapor S, Kapor S, Starcevic A, Popovic D, Djokovic A. Molecular Aspects of MAFLD-New Insights on Pathogenesis and Treatment. Curr Issues Mol Biol. 2023;45(11):9132–9148. doi:10.3390/cimb45110573

7. Donnelly KL, Smith CI, Schwarzenberg SJ, Jessurun J, Boldt MD, Parks EJ. Sources of fatty acids stored in liver and secreted via lipoproteins in patients with nonalcoholic fatty liver disease. J Clin Invest. 2005;115(5):1343–1351. doi:10.1172/JCI23621

8. Caron A, Richard D, Laplante M. The Roles of mTOR Complexes in Lipid Metabolism. Annu Rev Nutr. 2015;35:321–348. doi:10.1146/annurev-nutr-071714-034355

9. Mani S, Cao W, Wu L, Wang R. Hydrogen sulfide and the liver. Nitric Oxide Biol Chem. 2014;41:62–71. doi:10.1016/j.niox.2014.02.006

10. Yang G, Tang G, Zhang L, Wu L, Wang R. The pathogenic role of cystathionine γ-lyase/hydrogen sulfide in streptozotocin-induced diabetes in mice. Am J Pathol. 2011;179(2):869–879. doi:10.1016/j.ajpath.2011.04.028

11. Loiselle JJ, Yang G, Wu L. Hydrogen sulfide and hepatic lipid metabolism - a critical pairing for liver health. Br J Pharmacol. 2020;177(4):757–768. doi:10.1111/bph.14556

12. Kabil O, Motl N, Banerjee R. H2S and its role in redox signaling. Biochim Biophys Acta. 2014;1844(8):1355–1366. doi:10.1016/j.bbapap.2014.01.002

13. Whiteman M, Le Trionnaire S, Chopra M, Fox B, Whatmore J. Emerging role of hydrogen sulfide in health and disease: critical appraisal of biomarkers and pharmacological tools. Clin Sci Lond Engl 1979. 2011;121(11):459-488. doi:10.1042/CS20110267

14. Huang CW, Moore PK. H2S Synthesizing Enzymes: Biochemistry and Molecular Aspects. Handb Exp Pharmacol. 2015;230:3–25. doi:10.1007/978-3-319-18144-8_1

15. Nicholls P. Inhibition of cytochrome c oxidase by sulphide. Biochem Soc Trans. 1975;3(2):316–319.

16. Módis K, Coletta C, Erdélyi K, Papapetropoulos A, Szabo C. Intramitochondrial hydrogen sulfide production by 3-mercaptopyruvate sulfurtransferase maintains mitochondrial electron flow and supports cellular bioenergetics. FASEB J Off Publ Fed Am Soc Exp Biol. 2013;27(2):601–611. doi:10.1096/fj.12-216507

17. Módis K, Asimakopoulou A, Coletta C, Papapetropoulos A, Szabo C. Oxidative stress suppresses the cellular bioenergetic effect of the 3-mercaptopyruvate sulfurtransferase/hydrogen sulfide pathway. Biochem Biophys Res Commun. 2013;433(4):401–407. doi:10.1016/j.bbrc.2013.02.131

18. Szczesny B, Módis K, Yanagi K, Coletta C, Le Trionnaire S, Perry A, Wood ME, Whiteman M, Szabo C. AP39, a novel mitochondria-targeted hydrogen sulfide donor, stimulates cellular bioenergetics, exerts cytoprotective effects and protects against the loss of mitochondrial DNA integrity in oxidatively stressed endothelial cells in vitro. Nitric Oxide Biol Chem. 2014;41:120–130. doi:10.1016/j.niox.2014.04.008

19. Gerő D, Torregrossa R, Perry A, Waters A, Le-Trionnaire S, Whatmore JL, Wood M, Whiteman M. The novel mitochondria-targeted hydrogen sulfide (H2S) donors AP123 and AP39 protect against hyperglycemic injury in microvascular endothelial cells in vitro. Pharmacol Res. 2016;113(Pt A):186–198. doi:10.1016/j.phrs.2016.08.019

20. Karwi QG, Bornbaum J, Boengler K, Torregrossa R, Whiteman M, Wood ME, Schulz R, Baxter GF. AP39, a mitochondria-targeting hydrogen sulfide (H2 S) donor, protects against myocardial reperfusion injury independently of salvage kinase signalling. Br J Pharmacol. 2017;174(4):287–301. doi:10.1111/bph.13688

21. Trionnaire SL, Perry A, Szczesny B, Szabo C, Winyard PG, Whatmore JL, Wood ME, Whiteman M. The synthesis and functional evaluation of a mitochondria-targeted hydrogen sulfide donor, (10-oxo-10-(4-(3-thioxo-3H-1,2-dithiol-5-yl)phenoxy)decyl)triphenylphosphonium bromide (AP39). MedChemComm. 2014;5(6):728–736. doi:10.1039/C3MD00323J

22. Wiśniewski JR, Gaugaz FZ. Fast and sensitive total protein and Peptide assays for proteomic analysis. Anal Chem. 2015;87(8):4110–4116. doi:10.1021/ac504689z

23. Wiśniewski JR. Quantitative Evaluation of Filter Aided Sample Preparation (FASP) and Multienzyme Digestion FASP Protocols. Anal Chem. 2016;88(10):5438–5443. doi:10.1021/acs.analchem.6b00859

24. Bruderer R, Bernhardt OM, Gandhi T, Miladinović SM, Cheng LY, Messner S, Ehrenberger T, Zanotelli V, Butscheid Y, Escher C, Vitek O, Rinner O, Reiter L. Extending the limits of quantitative proteome profiling with data-independent acquisition and application to acetaminophen-treated three-dimensional liver microtissues. Mol Cell Proteomics MCP. 2015;14(5):1400–1410. doi:10.1074/mcp.M114.044305

25. Huang T, Bruderer R, Muntel J, Xuan Y, Vitek O, Reiter L. Combining Precursor and Fragment Information for Improved Detection of Differential Abundance in Data Independent Acquisition. Mol Cell Proteomics MCP. 2020;19(2):421–430. doi:10.1074/mcp.RA119.001705

26. Storey JD. A direct approach to false discovery rates. J R Stat Soc Ser B Stat Methodol. 2002;64(3):479–498. 10.1111/1467-9868.00346

27. Sundararaman N, Go J, Robinson AE, Mato JM, Lu SC, Van Eyk JE, Venkatraman V. PINE: An Automation Tool to Extract and Visualize Protein-Centric Functional Networks. J Am Soc Mass Spectrom. 2020;31(7):1410–1421. doi:10.1021/jasms.0c00032

28. Vizcaíno JA, Csordas A, Del-Toro N, Dianes JA, Griss J, Lavidas I, Mayer G, Perez-Riverol Y, Reisinger F, Ternent T, Xu QW, Wang R, Hermjakob H. 2016 update of the PRIDE database and its related tools. Nucleic Acids Res. 2016;44(D1):D447–456. doi:10.1093/nar/gkv1145

29. Whiteman M, Gooding KM, Whatmore JL, Ball CI, Mawson D, Skinner K, Tooke JE, Shore AC. Adiposity is a major determinant of plasma levels of the novel vasodilator hydrogen sulphide. Diabetologia. 2010;53(8):1722–1726. doi:10.1007/s00125-010-1761-5

30. Xu W, Cui C, Cui C, Chen Z, Zhang H, Cui Q, Xu G, Fan J, Han Y, Tang L, Targher G, Byrne CD, Zheng MH, Yang L, Cai J, Geng B. Hepatocellular cystathionine γ lyase/hydrogen sulfide attenuates nonalcoholic fatty liver disease by activating farnesoid X receptor. Hepatol Baltim Md. 2022;76(6):1794–1810. doi:10.1002/hep.32577

31. Li M, Xu C, Shi J, Ding J, Wan X, Chen D, Gao J, Li C, Zhang J, Lin Y, Tu Z, Kong X, Li Y, Yu C. Fatty acids promote fatty liver disease via the dysregulation of 3-mercaptopyruvate sulfurtransferase/hydrogen sulfide pathway. Gut. 2018;67(12):2169–2180. doi:10.1136/gutjnl-2017-313778

32. Sun L, Zhang S, Yu C, Pan Z, Liu Y, Zhao J, Wang X, Yun F, Zhao H, Yan S, Yuan Y, Wang D, Ding X, Liu G, Li W, Zhao X, Liu Z, Li Y. Hydrogen sulfide reduces serum triglyceride by activating liver autophagy via the AMPK-mTOR pathway. Am J Physiol Endocrinol Metab. 2015;309(11):E925–935. doi:10.1152/ajpendo.00294.2015

33. Wu D, Zhong P, Wang Y, Zhang Q, Li J, Liu Z, Ji A, Li Y. Hydrogen Sulfide Attenuates High-Fat Diet-Induced Non-Alcoholic Fatty Liver Disease by Inhibiting Apoptosis and Promoting Autophagy via Reactive Oxygen Species/Phosphatidylinositol 3-Kinase/AKT/Mammalian Target of Rapamycin Signaling Pathway. Front Pharmacol. 2020;11:585860. doi:10.3389/fphar.2020.585860

34. Wu D, Zheng N, Qi K, Cheng H, Sun Z, Gao B, Zhang Y, Pang W, Huangfu C, Ji S, Xue M, Ji A, Li Y. Exogenous hydrogen sulfide mitigates the fatty liver in obese mice through improving lipid metabolism and antioxidant potential. Med Gas Res. 2015;5(1):1. doi:10.1186/s13618-014-0022-y

35. Cui X, Yao M, Feng Y, Li C, Li Y, Guo D, He S. Exogenous hydrogen sulfide alleviates hepatic endoplasmic reticulum stress via SIRT1/FoxO1/PCSK9 pathway in NAFLD. FASEB J Off Publ Fed Am Soc Exp Biol. 2023;37(8):e23027. doi:10.1096/fj.202201705RR

36. Shimano H, Sato R. SREBP-regulated lipid metabolism: convergent physiology - divergent pathophysiology. Nat Rev Endocrinol. 2017;13(12):710–730. doi:10.1038/nrendo.2017.91

37. Moon YA. The SCAP/SREBP Pathway: A Mediator of Hepatic Steatosis. Endocrinol Metab Seoul Korea. 2017;32(1):6–10. doi:10.3803/EnM.2017.32.1.6

38. Zhang S, Wang M, Li H, Li Q, Liu N, Dong S, Zhao Y, Pang K, Huang J, Ren C, Wang Y, Tian Z, Lu F, Zhang W. Exogenous H2 S promotes ubiquitin-mediated degradation of SREBP1 to alleviate diabetic cardiomyopathy via SYVN1 S-sulfhydration. J Cachexia Sarcopenia Muscle. 2023;14(6):2719–2732. doi:10.1002/jcsm.13347

39. Sun Q, Xing X, Wang H, Wan K, Fan R, Liu C, Wang Y, Wu W, Wang Y, Wang R. SCD1 is the critical signaling hub to mediate metabolic diseases: Mechanism and the development of its inhibitors. Biomed Pharmacother Biomedecine Pharmacother. 2023;170:115586. doi:10.1016/j.biopha.2023.115586

40. Miyazaki M, Flowers MT, Sampath H, Chu K, Otzelberger C, Liu X, Ntambi JM. Hepatic stearoyl-CoA desaturase-1 deficiency protects mice from carbohydrate-induced adiposity and hepatic steatosis. Cell Metab. 2007;6(6):484–496. doi:10.1016/j.cmet.2007.10.014

41. Ntambi JM, Miyazaki M, Stoehr JP, Lan H, Kendziorski CM, Yandell BS, Song Y, Cohen P, Friedman JM, Attie AD. Loss of stearoyl-CoA desaturase-1 function protects mice against adiposity. Proc Natl Acad Sci U S A. 2002;99(17):11482–11486. doi:10.1073/pnas.132384699

42. Wang Y, Nakajima T, Gonzalez FJ, Tanaka N. PPARs as Metabolic Regulators in the Liver: Lessons from Liver-Specific PPAR-Null Mice. Int J Mol Sci. 2020;21(6):2061. doi:10.3390/ijms21062061

43. Inoue J, Yamasaki K, Ikeuchi E, Satoh S ichi, Fujiwara Y, Nishimaki-Mogami T, Shimizu M, Sato R. Identification of MIG12 as a mediator for stimulation of lipogenesis by LXR activation. Mol Endocrinol Baltim Md. 2011;25(6):995–1005. doi:10.1210/me.2011-0070

44. Singh R, Wang Y, Xiang Y, Tanaka KE, Gaarde WA, Czaja MJ. Differential effects of JNK1 and JNK2 inhibition on murine steatohepatitis and insulin resistance. Hepatol Baltim Md. 2009;49(1):87–96. doi:10.1002/hep.22578

45. Avruch J, Lin Y, Long X, Murthy S, Ortiz-Vega S. Recent advances in the regulation of the TOR pathway by insulin and nutrients. Curr Opin Clin Nutr Metab Care. 2005;8(1):67–72. doi:10.1097/00075197-200501000-00010

46. Wheatcroft SB, Kearney MT, Shah AM, Ezzat VA, Miell JR, Modo M, Williams SCR, Cawthorn WP, Medina-Gomez G, Vidal-Puig A, Sethi JK, Crossey PA. IGF-binding protein-2 protects against the development of obesity and insulin resistance. Diabetes. 2007;56(2):285–294. doi:10.2337/db06-0436

47. Dan HC, Cooper MJ, Cogswell PC, Duncan JA, Ting JPY, Baldwin AS. Akt-dependent regulation of NF-κB is controlled by mTOR and Raptor in association with IKK. Genes Dev. 2008;22(11):1490–1500. doi:10.1101/gad.1662308

48. Zhao M, Cheng Y, Wang X, Cui X, Cheng X, Fu Q, Song Y, Yu P, Liu Y, Yu Y. Hydrogen Sulfide Attenuates High-Fat Diet-Induced Obesity: Involvement of mTOR/IKK/NF-κB Signaling Pathway. Mol Neurobiol. 2022;59(11):6903–6917. doi:10.1007/s12035-022-03004-0

49. Ghosh S, Hayden MS. New regulators of NF-kappaB in inflammation. Nat Rev Immunol. 2008;8(11):837–848. doi:10.1038/nri2423

50. Jiang B, Wang D, Hu Y, Li W, Liu F, Zhu X, Li X, Zhang H, Bai H, Yang Q, Yang X, Ben J, Chen Q. Serum amyloid A1 exacerbates hepatic steatosis via TLR4-mediated NF-κB signaling pathway. Mol Metab. 2022;59:101462. doi:10.1016/j.molmet.2022.101462

51. Wang Y, Cao F, Wang Y, Yu G, Jia BL. Silencing of SAA1 inhibits palmitate- or high-fat diet induced insulin resistance through suppression of the NF-κB pathway. Mol Med Camb Mass. 2019;25(1):17. doi:10.1186/s10020-019-0075-4

52. Kanamori M, Suzuki H, Saito R, Muramatsu M, Hayashizaki Y. T2BP, a novel TRAF2 binding protein, can activate NF-kappaB and AP-1 without TNF stimulation. Biochem Biophys Res Commun. 2002;290(3):1108–1113. doi:10.1006/bbrc.2001.6315

53. Okumura A, Unoki-Kubota H, Matsushita Y, Shiga T, Moriyoshi Y, Yamagoe S, Kaburagi Y. Increased serum leukocyte cell-derived chemotaxin 2 (LECT2) levels in obesity and fatty liver. Biosci Trends. 2013;7(6):276–283.

54. Takata N, Ishii KA, Takayama H, Nagashimada M, Kamoshita K, Tanaka T, Kikuchi A, Takeshita Y, Matsumoto Y, Ota T, Yamamoto Y, Yamagoe S, Seki A, Sakai Y, Kaneko S, Takamura T. LECT2 as a hepatokine links liver steatosis to inflammation via activating tissue macrophages in NASH. Sci Rep. 2021;11(1):555. doi:10.1038/s41598-020-80689-0

55. Choi YJ, Kim S, Choi Y, Nielsen TB, Yan J, Lu A, Ruan J, Lee HR, Wu H, Spellberg B, Jung JU. SERPINB1-mediated checkpoint of inflammatory caspase activation. Nat Immunol. 2019;20(3):276–287. doi:10.1038/s41590-018-0303-z

56. Kamal MM, Adel A, Sayed GH, Ragab S, Kassem DH. New emerging roles of the novel hepatokine SERPINB1 in type 2 diabetes mellitus: Crosstalk with β-cell dysfunction and dyslipidemia. Transl Res J Lab Clin Med. 2021;231:1–12. doi:10.1016/j.trsl.2020.12.004

57. Zhao S, Li N, Xiong W, Li G, He S, Zhang Z, Zhu Q, Jiang N, Ikejiofor C, Zhu Y, Wang MY, Han X, Zhang N, Solis-Herrera C, Kusminski C, An Z, Elmquist JK, Scherer PE. Leptin Reduction as a Required Component for Weight Loss. Diabetes. 2023;73(2):197–210. doi:10.2337/db23-0571

58. Goran MI, Alderete TL. Targeting Adipose Tissue Inflammation to Treat the Underlying Basis of the Metabolic Complications of Obesity. Published online October 26, 2012. doi:10.1159/000341287

59. Ojha S, Budge H, Symonds ME. Adipocytes in Normal Tissue Biology. In: McManus LM, Mitchell RN, eds. Pathobiology of Human Disease. Academic Press; 2014:2003–2013. doi:10.1016/B978-0-12-386456-7.04408-7

60. Mao S, Wang X, Li M, Liu H, Liang H. The role and mechanism of hydrogen sulfide in liver fibrosis. Nitric Oxide Biol Chem. Published online February 13, 2024:S1089–8603(24)00019-3. doi:10.1016/j.niox.2024.02.002

61. Lohakul J, Jeayeng S, Chaiprasongsuk A, Torregrossa R, Wood MarkE, Saelim M, Thangboonjit W, Whiteman M, Panich U. Mitochondria-Targeted Hydrogen Sulfide Delivery Molecules Protect Against UVA-Induced Photoaging in Human Dermal Fibroblasts, and in Mouse Skin In Vivo. Antioxid Redox Signal. 2022;36(16-18):1268–1288. doi:10.1089/ars.2020.8255

