## Supplemental Figures for "Mitochondria-targeted hydrogen sulfide donor reduces fatty liver and obesity in mice fed a high fat diet by inhibiting *de novo* lipogenesis and inflammation *via* mTOR/SREBP-1 and NF-κB signaling pathways"

**Supplemental Figure 1**

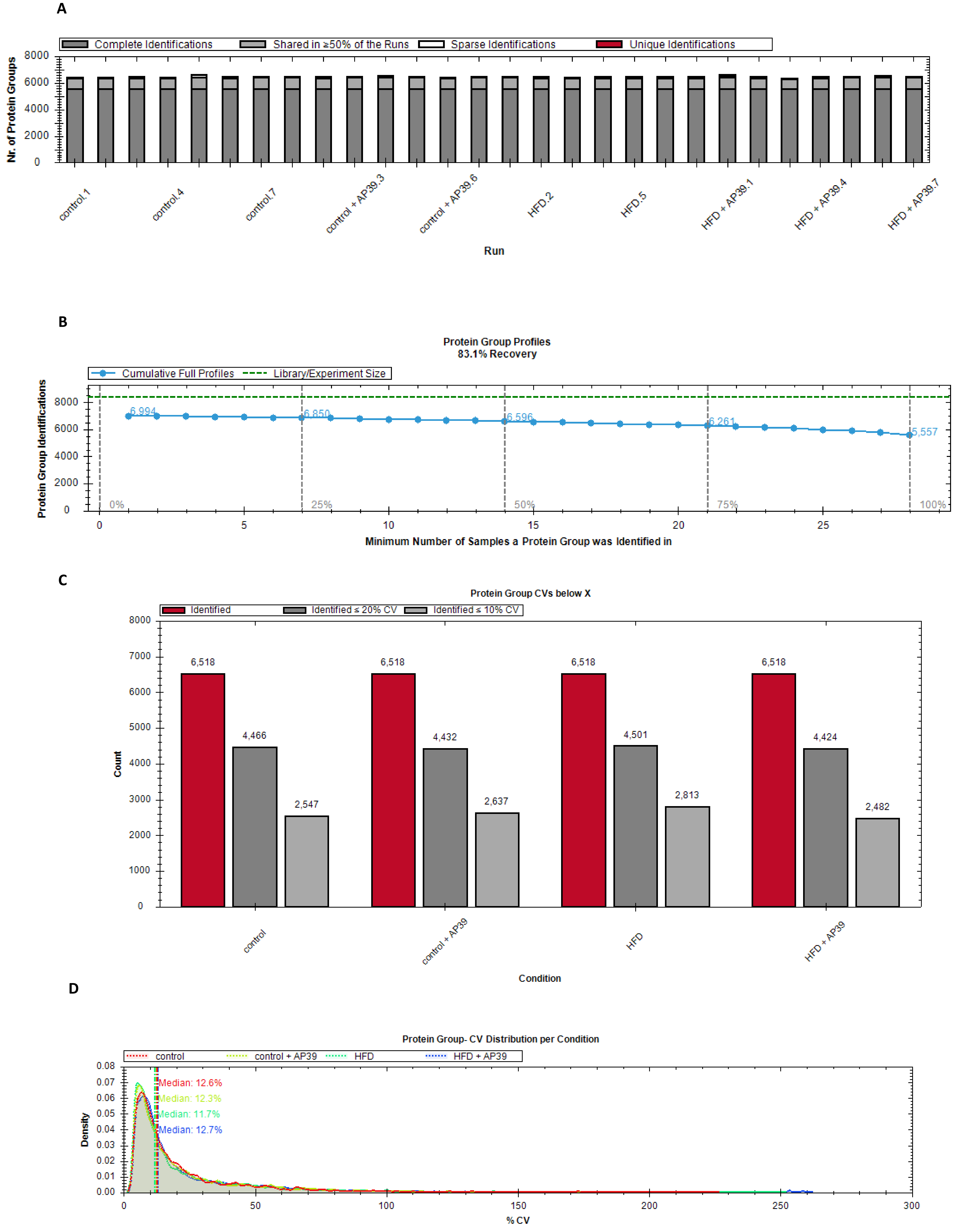

E

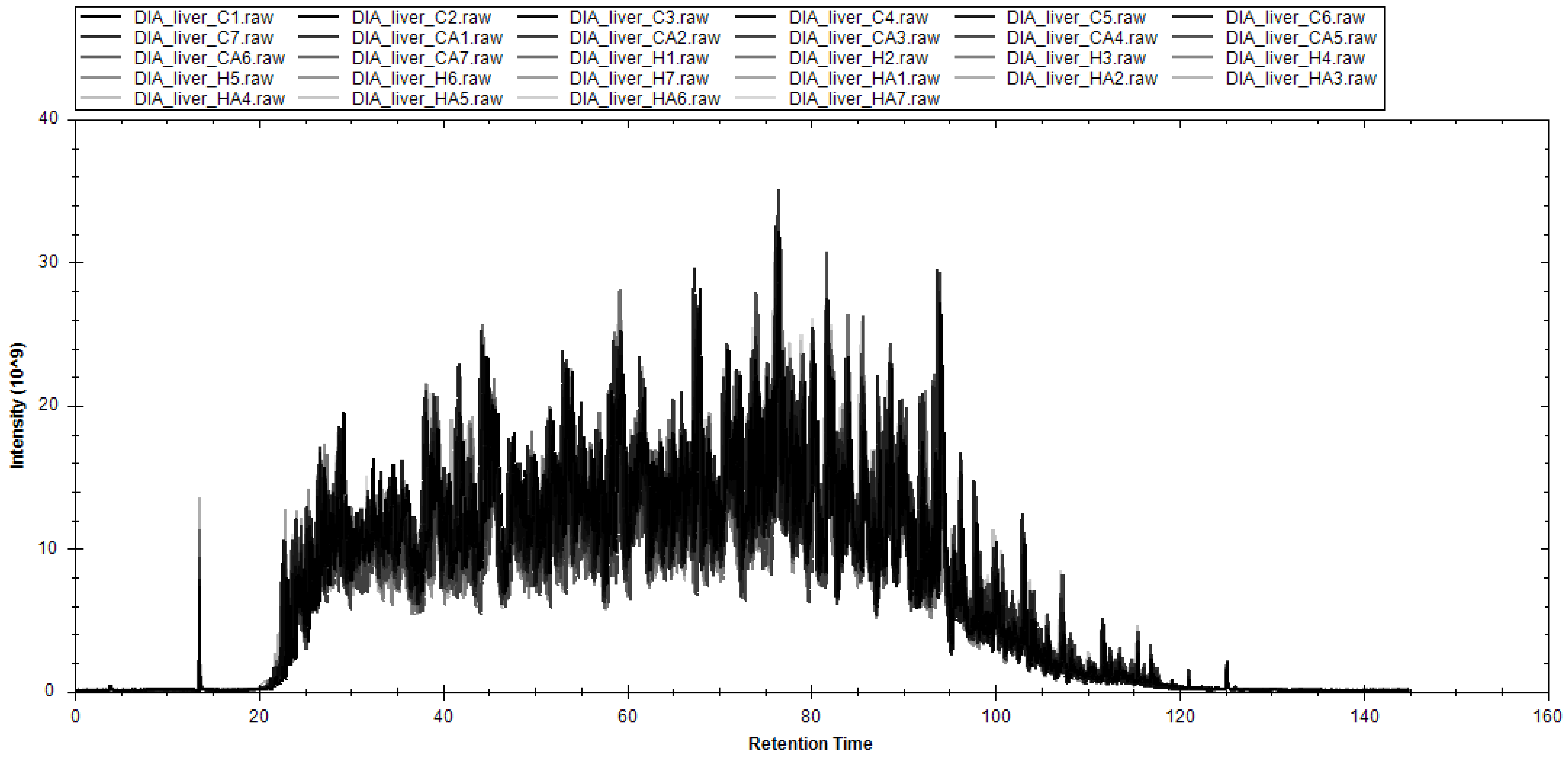
DIA_liver_C1.raw
DIA_liver_C2.raw
DIA_liver_C3.raw
DIA_liver_C4.raw
DIA_liver_C5.raw
DIA_liver_C6.raw
DIA_liver_C7.raw
DIA_liver_CA1.raw
DIA_liver_CA2.raw
DIA_liver_CA3.raw
DIA_liver_CA4.raw
DIA_liver_CA5.raw
DIA_liver_CA6.raw
DIA_liver_CA7.raw
DIA_liver_H1.raw
DIA_liver_H2.raw
DIA_liver_H3.raw
DIA_liver_H4.raw
DIA_liver_H5.raw
DIA_liver_H6.raw
DIA_liver_H7.raw
DIA_liver_HA1.raw
DIA_liver_HA2.raw
DIA_liver_HA3.raw
DIA_liver_HA4.raw
DIA_liver_HA5.raw
DIA_liver_HA6.raw
DIA_liver_HA7.raw
Intensity (10^9)
Retention Time
0
10
20
30
40
0
20
40
60
80
100
120
140
160

F

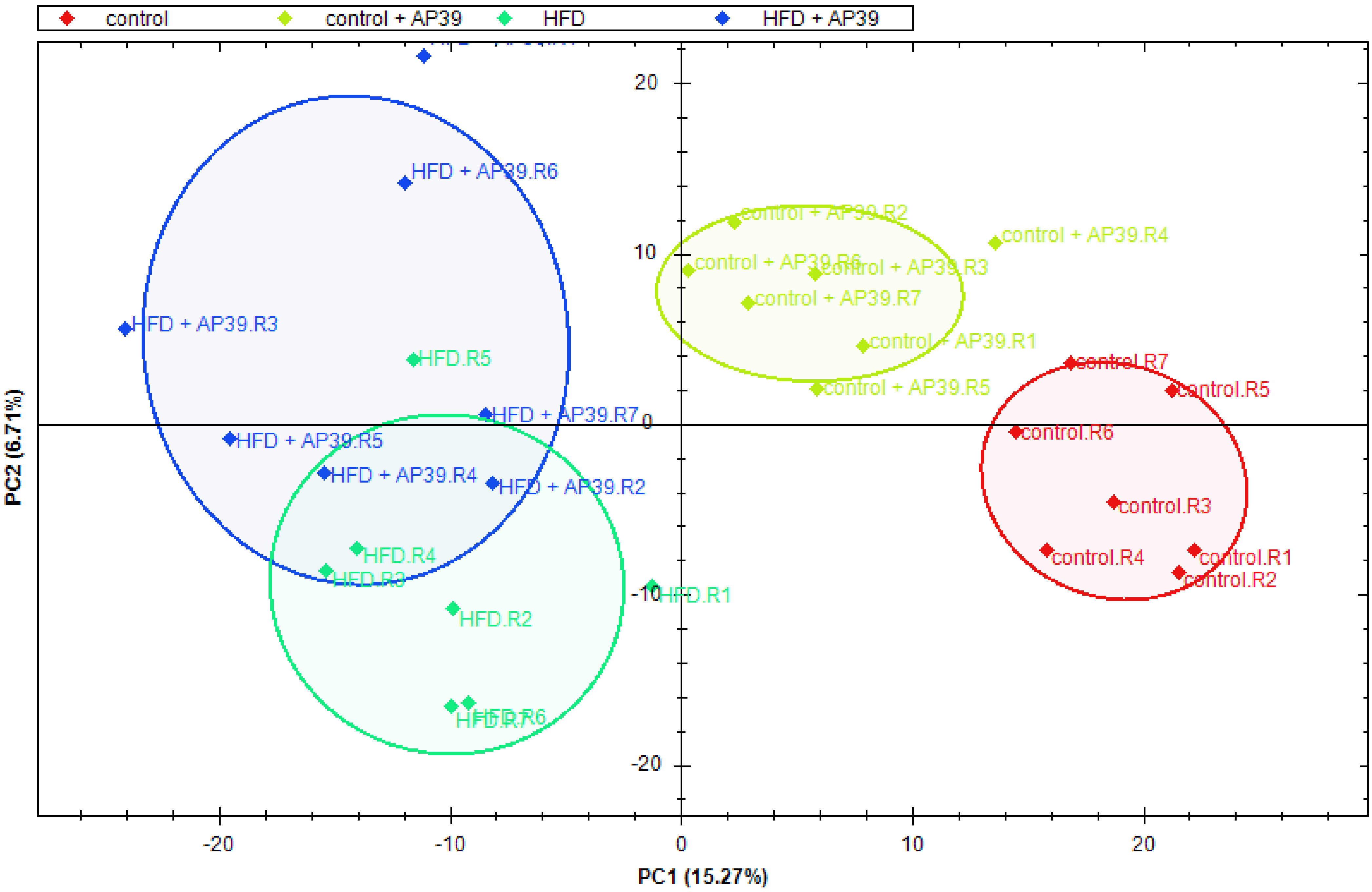
control
control + AP39
HFD
HFD + AP39
PC2 (6.71%)
PC1 (15.27%)
HFD + AP39.R6
HFD + AP39.R3
HFD.R5
HFD + AP39.R7
HFD + AP39.R5
HFD + AP39.R4
HFD + AP39.R2
HFD.R4
HFD.R3
HFD.R2
HFD.R1
HFD.R6
HFD.R7
control + AP39.R2
control + AP39.R4
control + AP39.R6
control + AP39.R3
control + AP39.R7
control + AP39.R1
control + AP39.R5
control.R7
control.R5
control.R6
control.R3
control.R4
control.R1
control.R2
20
10
0
-10
-20
-20
-10
0
10
20

Supplemental Figure 2

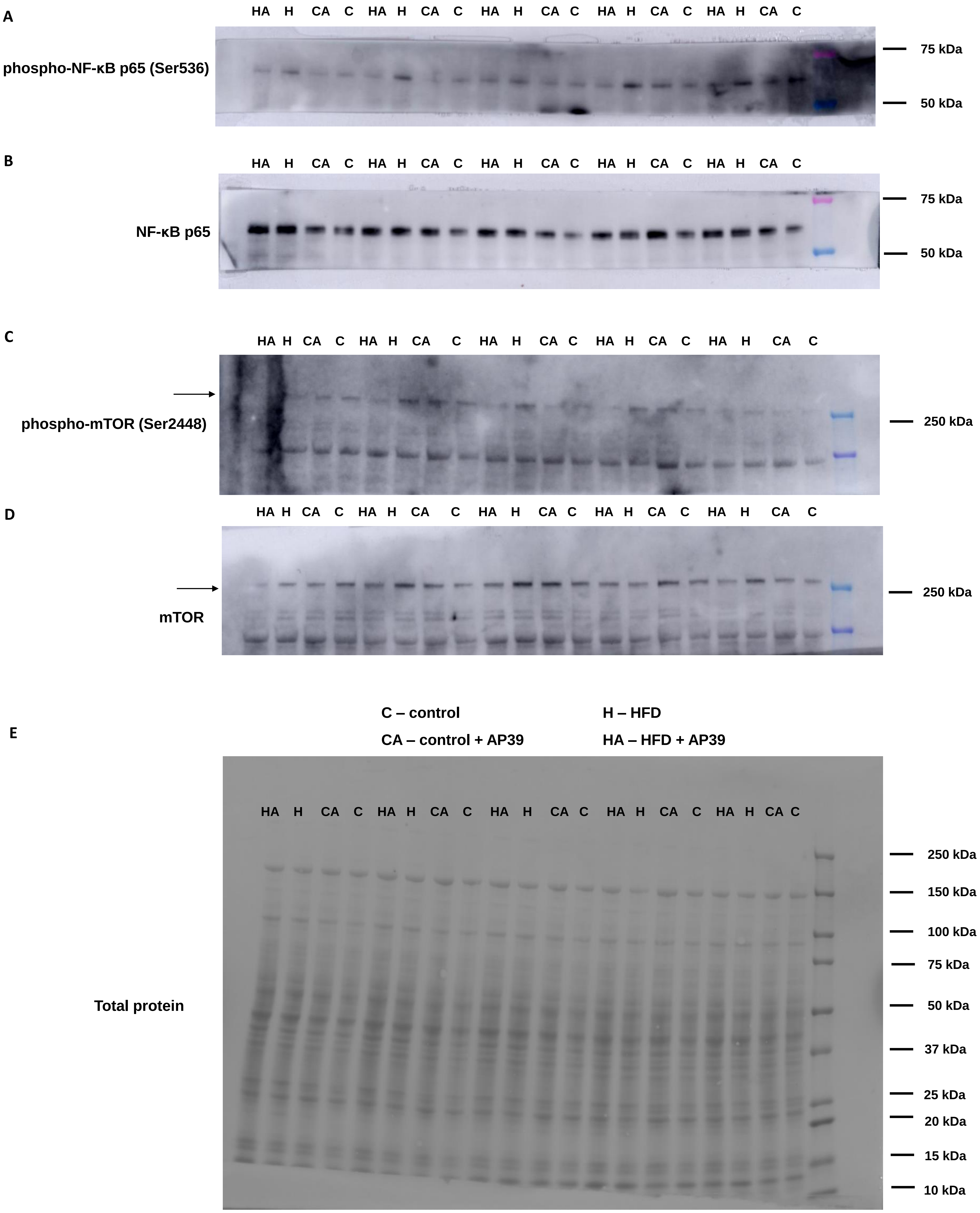

**Supplemental Figure 3**

**A**

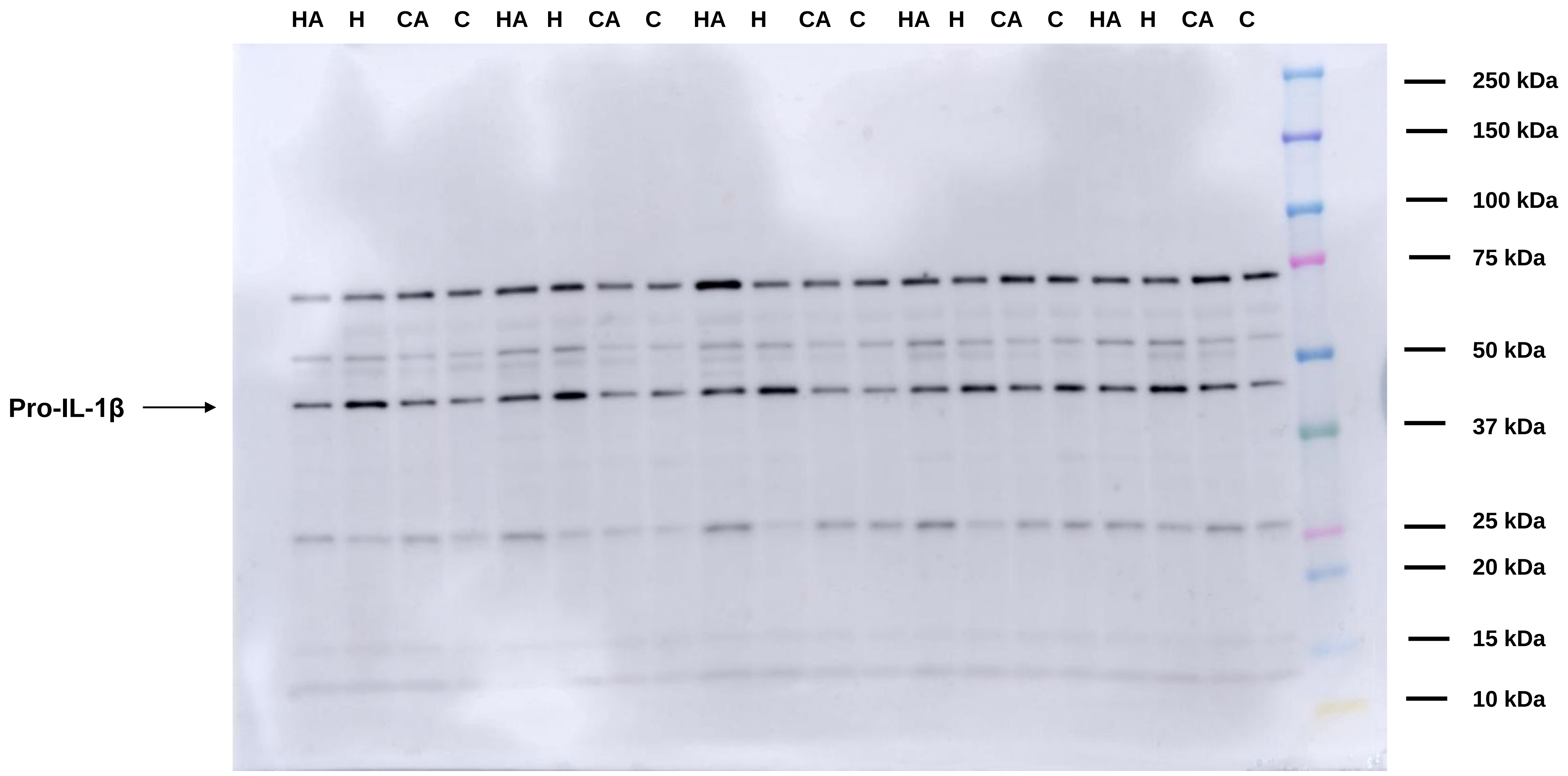


C – control

CA – control + AP39

H – HFD

HA – HFD + AP39

**B**

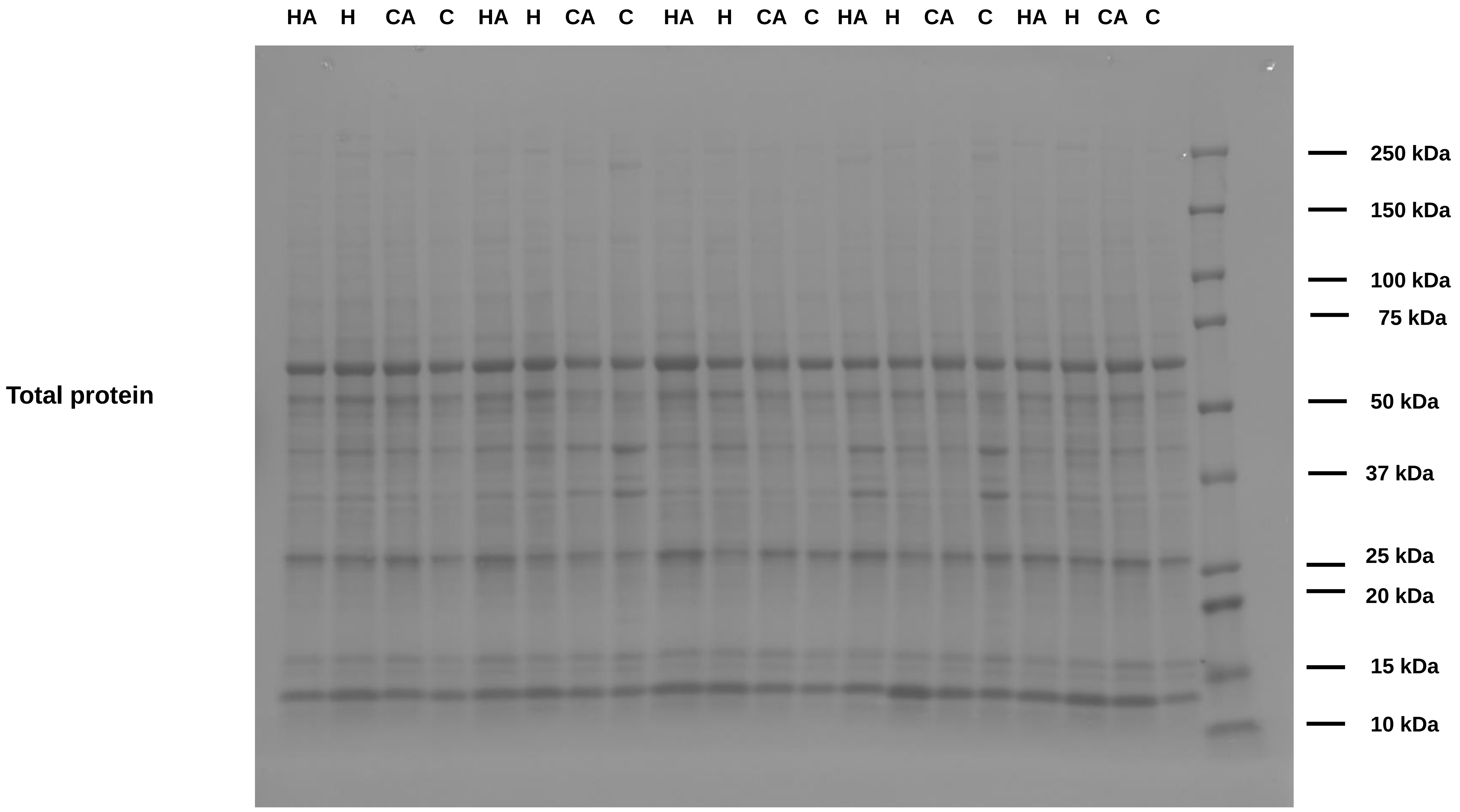

**<u>Supplemental Figure 1</u>** **Quality control of MS runs of the liver of control, control + AP39, high fat diet (HFD) and HFD + AP39 groups.** Protein group identification details across all LC-MS runs **(A)**. Spectral library recovery **(B)**. Coefficient of variations (CVs) for protein groups across all biological conditions **(C)**. Distribution of protein group CV in biological conditions **(D)**. Total ion chromatogram (TIC) overlay of all LC-MS runs **(E)**. Principal component analysis of the proteomic data of the liver of control, control + AP39, HFD and HFD + AP39 groups in a 2D graph of principal component 1 and component 2.

**<u>Supplemental Figure 2</u>** **Western blot replicates as uncropped images. Related to Figure 5M.** Western blot replicates for phospho-NF-κB (nuclear factor kappa-light-chain-enhancer of activated B cells) p65 (Ser536) **(A)**, NF-κB p65 **(B)**, phospho-mTOR (mammalian target of rapamycin) (Ser2448) **(C)**, mTOR **(D)** in the liver across all biological conditions. Ponceau S staining used to normalize data for total protein level **(E)** (n=5).

**<u>Supplemental Figure 3</u>** **Western blot replicates as uncropped images. Related to Figure 7L.** Western blot replicates for pro-IL-1β (pro-interleukin 1 beta) **(A)** in the eWAT across all biological conditions. Ponceau S staining used to normalize data for total protein level **(B)** (n=5).
